# Integrating functional genomics and proteomics identifies Folate Carrier SLC19A1 as a predictor of pralatrexate sensitivity in T-cell lymphoma

**DOI:** 10.1101/2025.10.08.681217

**Authors:** Jacob C. Pantazis, Amy E. Pomeroy, Adam C. Palmer

## Abstract

Cancer therapies are typically effective in subsets of patients, reflecting the molecular diversity of cancers and motivating the need for predictive biomarkers of response. Biomarker-guided therapy is increasingly useful in oncology, yet biomarker discovery remains complicated by the large number of molecular features that make it difficult to distinguish causal determinants from spurious associations. To address this challenge, we combined functional genomic screening, proteomics, and drug sensitivity profiling to discover response biomarkers for a number of therapies used in the treatment of Peripheral T-Cell Lymphomas (PTCL). First, we used genome-wide CRISPR-dCas9 interference screens in PTCL cells under drug treatment to identify a shortlist of genes whose knockdown directly increases or decreases drug sensitivity. Next, we profiled drug responses across a diverse panel of 30 PTCL cultures and, from the shortlist, identified genes whose protein abundance correlated with drug sensitivity. Genes detected by both approaches are causal determinants of drug response and correlates of drug response across the panel of lymphoma cultures, making them promising candidates for predictive biomarkers. Basal expression of the reduced folate carrier SLC19A1 was a strong predictor of pralatrexate sensitivity, consistent with its role as the primary transporter for pralatrexate uptake. Simulated clinical trials predicted that biomarker-guided patient selection could improve the power to detect significant benefit of adding pralatrexate to frontline chemotherapy in PTCL. These findings illustrate how the causal insights of functional genetic screens can augment correlative studies to identify biomarkers of drug response, and suggest the potential for precise use of pralatrexate for PTCL.

## INTRODUCTION

Most cancer therapies are effective in only a subset of patients, reflecting molecular heterogeneity among patients’ cancers. A proven strategy to improve the success of oncology trials is to use biomarkers to identify the patient population in which a particular therapy is most effective^1^. For some therapies, target sequence or expression can be an effective biomarker, such as inhibitors of mutant or fusion oncogenes (*EGFR*, *BRAF*, *ALK*, *BCR-ABL*)^2–5^ and antibody-based therapies (HER2, PD-L1, CD30)^6–8^. However, most cancer therapies lack obvious biomarkers, and determinants of drug sensitivity can extend beyond the target to include processes such as cellular entry, efflux, metabolism, pro-drug activation, and regulation of the target pathway^9–11^. For some therapies, response may be so polygenic as to make prediction impractical, yet clinical precedent shows that for some drugs the status of one or a few genes (other than the target) is sufficient to identify responsive patient populations, such as *KRAS* for the anti-EGFR antibody cetuximab, and *BRCA* for PARP inhibition^12,13^. Similarly informative biomarkers might be found for existing cancer therapies.

Pharmacogenomic resources such as the Cancer Therapeutics Response Portal (CTRP) and Genomics of Drug Sensitivity in Cancer (GDSC) have advanced our understanding of drug mechanisms by profiling drug responses across panels of cancer cell lines and identifying molecular features, such as gene expression or mutations, that correlate with sensitivity to particular drugs^10,14^. Such data provide mechanistic insight and rich hypotheses for biomarker discovery, but searching for correlations among over ten thousand genes typically yields hundreds to thousands of nominally significant correlates per drug. Potentially useful biomarkers may lie in these data but are difficult to distinguish from indirect or spurious correlations. Combining prior knowledge with chemical or genetic knockdown of individual genes has proven a useful approach to confirm determinants of drug activity^15^, but bespoke approaches are difficult to scale. Pooled CRISPR-Cas9 knockdown screens, when performed under drug treatment, offer an unbiased and genome-scale approach to assess the influence of individual genes upon drug resistance or hypersensitivity in an isogenic background^16,17^. Here we combine such functional genetic screens with correlative analyses to identify predictive biomarkers, where knockdown screens narrow the search space to genes with proven causal impact on drug response, and correlative studies across diverse models then identify which of these also predict sensitivity. Thus, orthogonal criteria – causal and correlative – enrich for meaningful biomarkers of drug response, analogous to finding a needle in a haystack using dual search criteria of ‘in the haystack’ and ‘made of metal’.

Aggressive lymphomas are widely recognized as promising candidates for precision medicine^18^, making them a compelling setting to test the integration of correlative and causal methods for biomarker discovery. Precision strategies are desirable in aggressive lymphomas, such as Peripheral T-Cell Lymphomas (PTCL), because (1) several clinically effective therapies are available to choose from^19^, (2) most therapies are effective in a minority of patients^20–22^, (3) most therapies presently lack biomarkers to guide selection^23^, and (4) when biomarkers are known, precision combination therapies have improved cure rates^8^. Recent clinical trials highlight the promise and challenges of precision strategies. The PHOENIX trial of ibrutinib plus R-CHOP in diffuse large B-cell lymphoma (DLBCL) was negative overall, but improved survival in a genetically defined subset demonstrated that testing an effective therapy in a broad population can obscure its benefits^24^. The phase 2 GUIDANCE-01 trial showed improved survival by assigning each of six DLBCL subtypes to targeted agents plus RCHOP^25^. However, the assignment of drugs to subtypes in GUIDANCE-01 was largely based on intuition and it is unclear which assignments are accurate; the success of this approach could be aided by evidence-based biomarkers. In Peripheral T-cell Lymphomas (PTCL), the ECHELON-2 trial showed that CHP plus brentuximab vedotin, an antibody-drug conjugate targeting CD30, significantly improved cure rates and 5-year overall survival in CD30+ PTCL^8^, demonstrating the potential of biomarker-guided combinations when predictive biomarkers are known.

In this study we aimed to discover predictive biomarkers of drug response in Peripheral T-cell Lymphomas (PTCL). PTCLs are particularly heterogeneous because the immune system comprises many types of T-lymphocyte that can become malignant in many ways^26,27^. First-line combination regimens including CHOP, CHOEP, and CH(E)P-brentuximab vedotin cure some patients, but survival is poor outside of the ALK-positive anaplastic large cell lymphoma (ALCL) subtype. A variety of drug classes have demonstrated clinical efficacy against relapsed or refractory (r/r) disease, including DNA-damaging chemotherapies gemcitabine and oxaliplatin, histone deacetylase inhibitors romidepsin and belinostat, the proteasome inhibitor bortezomib, and the folate synthesis inhibitor pralatrexate^20–22,28^. These therapies can induce complete responses in approximately 20 to 30% of patients, showing their potential to improve outcomes for some patients, yet lacking the broad activity that is often required to substantially improve front-line combination therapy for all patients. Although histone deacetylase inhibitors are most effective against subtypes known as T Follicular Helper lymphomas^29^, the activity of other available therapies are not well delineated by histologic subtype, establishing an unmet need for biomarker discovery in these diseases.

To discover drug response biomarkers for agents with clinical activity in relapsed/refractory PTCL, here we combined orthogonal experimental methods that provide independent information on causality and correlation between gene expression and drug sensitivity. Genome-wide CRISPR interference screens under drug treatment identified genes whose knockdown caused significant drug resistance or hypersensitivity. We then profiled drug responses and proteomics across a panel of 30 mature T– and NK-cell lymphoma cultures, and analyzed correlations between protein levels and drug sensitivity. For each drug, 2 to 5 genes were concordant hits by both methods, from which we developed simple protein expression signatures (sum of z-scores) and assessed how accurately these potential biomarkers distinguish sensitive from resistant lymphoma cultures. The folate carrier SLC19A1 stood out as the top single-gene predictor of pralatrexate sensitivity in PTCL, consistent with its role as the principal transporter facilitating cellular uptake of reduced folates as well as antifolates pralatrexate and methotrexate^30^. Using the measured accuracy of SLC19A1 in predicting pralatrexate sensitivity, we simulated combination therapy trials with and without biomarker selection, which showed that biomarker-guided therapy improves the power to significantly improve survival in frontline PTCL.

## MATERIALS AND METHODS

### Cell culture

T-cell and Natural Killer-cell lymphoma cell lines were procured from the American Type Culture Collection (ATCC), JCRB Cell Bank, Leibniz Institute DSMZ, and Sigma-Aldrich by the lab of David Weinstock before use in this study^31^. Three additional Peripheral T Cell Lymphoma Cultures (OCI-LY12, OCI-LY13.2, and OCI-LY17, RRID’s: CVCL_8796, CVCL_8797, and CVCL_8798 respectively) were purchased from the University Health Network in Toronto, ON. The identities of cell cultures were confirmed by STR profiling performed by Genetica Cell Line Services (RRID: SCR_014504) using 20ng samples of genomic DNA, extracted from each culture with QIAGEN DNeasy Blood and Tissue kits (Catalog number: 69504) (**Supplementary Table 1**).

All cell lines were maintained in Human Plasma-Like Medium (Gibco A4899101). This media was supplemented with 10% (v/v) heat inactivated fetal bovine serum (FBS) (Gibco 16140071), 2 mM L-alanine-L-glutamine (GlutaMAX), and penicillin/streptomycin 100 U/mL and 100 μg/mL (Gibco). Additionally, some cultures were dependent upon human recombinant IL-2 (Peprotech 200-02), supplemented at a final concentration of 100 U/mL. Cultures were grown at 37°C and 5% CO2. Cultures were tested for mycoplasma contaminants following seeding from frozen aliquots and alongside drug response experiments (Lonza MycoAlert). Live cell counts were monitored every 48-72 hours with trypan blue with passaging by establishing fresh cultures at a consistent seeding density of 100,000 live cells per mL.

### PTCL therapies

Five therapies used in the treatment of T-cell Lymphoma were purchased from MedChemExpress; romidepsin (Catalog number: HY15149, pralatrexate (Catalog number: HY-10466), bortezomib (Catalog Number: HY-10227), gemcitabine (Catalog Number: HY-17026), oxaliplatin (Catalog number: HY-17371). Stock solutions were prepared in DMSO 10mM and stored at –80C, except oxaliplatin which was stored in phosphate buffered saline to maintain drug activity. For drug sensitivity profiling, therapies were thawed and diluted to working concentrations prior to dose response measurements. Dose response ranges were centered around average blood plasma concentrations of each therapy 12 hours after drug administration, identified from human pharmacokinetic studies^32–36^.

### Dose response measurements

Prior to drug treatment, all T and NK cultures were passaged every 48 hours to an initial concentration of 100,000 live cells/mL for approximately a week, establishing consistent exponential growth. A target of 3500 live cells in 35 μL of complete HPLM were dispensed into each well of black 384 well plates (ThermoFisher Multidrop Combi, RRID: SCR_019329). A D300e Digital Drug Dispenser (Tecan, RRID: SCR_027242) was used to dispense 16-point concentration gradients of each therapy across each plate. Well locations were randomized by D300 eControl software to avoid systematic error across plates, and data were ‘descrambled after measurements for analysis. Each plate contained a minimum of 96 untreated control wells for normalization. These plates were then incubated at 37 °C in 5% CO_2_ for 72 hours in humidity-controlled boxes. At 72 hours, 35 μL of CellTiterGlo 2.0 (Promega Catalog Number: G9243) diluted 1:1 in phosphate-buffered saline was added to each well plate. Plates were incubated at room temperature for 20 minutes, and luminescence was measured using a BMG ClarioSTAR^plus^ (RRID: SCR_026330), which was the plate reader of choice for these experiments due to its automated cross-talk reduction between adjacent wells. For rigor and reproducibility, these dose-response measurements were repeated in parallel cultures that had been passaged separately up until the collection of dose responses. These measurements were repeated on replicate weeks in these parallel cultures (**Supplementary Figures 1 and 2**). Hill functions were fitted to each dose response according to the model

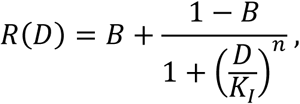

where *D* is drug dose, and *B*, *K_I_*, and *n* are the baseline, the half-maximal inhibitory concentration, and Hill coefficient respectively (implemented by NonlinearModelFit in Mathematica 13). Drug sensitivity was quantified by calculating the area over the curve (AOC) for each fitted response on a log_10_ scale, since the area ‘under’ a curve on a log scale is infinite. Alongside dose response measurements, the growth of untreated cells was monitored in 12 well culture plates at a consistent surface area to volume ratio compared to 384 well plate formats to observe consistency in doubling rate for cell cultures (**Supplementary Figure 3**).

### Production of dCas9-expressing PTCL cultures

Lentiviral vectors containing dCas9 constructs were produced in HEK293T cells (RRID: CVCL_0063, obtained from the UNC Tissue Culture Facility, RRID: SCR_012432). HEK293T cells, grown in 10cm^2^ plates, were transfected with the pHR-SFFV-dCas9-BFP-KRAB plasmid (RRID: Addgene_46911) along with Lentiviral packaging plasmids psPAX2 (RRID: Addgene_12260) and pMD2.G (RRID: Addgene_12259) using 6 μg/mL of 2500 Molecular weight polyethylenimine (Fisher 50-255-9824). HEK293T cells were maintained in complete Dulbecco’s Medium plus 20mM HEPES during transfection and grown for three days in ten 10 cm^2^ dishes at 37 °C, 5% CO_2_. Lentivirus containing the dCas9-BFP-KRAB insert was harvested at 48 and 72 hours. Lentiviral harvests were sterile filtered and concentrated by centrifuge (3350x g at 4 °C overnight) using LentiFuge viral concentration reagent (Cellecta catalog number: LFVC1). Pelleted lentivirus was resuspended in chilled, sterile PBS and stored at –80C.

Two million SU-DHL-1 (DMSZ Catalog Number, ACC356, RRID: CVCL_0538) cells were infected with dCas9-TagBFP-KRAB lentivirus in 1mL of infection media. Infection media contained RPMI-1640 (Gibco) plus 20mM HEPES and 8 ug/mL polybrene (Sigma TR-1003). Cells were centrifuged in infection media for 1 hour at 1000x g, then returned to 10mL of complete RPMI-1640. After 7-10 days of growth, the proportion of infected cells – which express Blue Fluorescent Protein (TagBFP) – was measured by an Attune NxT flow cytometer (RRID: SCR_0919590) using the violet laser (405/457 Ex/Em). The population of infected cells was enriched to greater than 90% with three rounds of cell sorting a FACS Aria II (RRID: SCR_018934) by the UNC Flow Cytometry Core Facility (RRID:SCR_019170).

Confirmation of dCas9-TagBFP-KRAB activity was performed with the FACS-sorted culture by infecting cells with lentivirus-packaged guide RNAs. 7 to 10 days after expansion and selection of guide RNA-positive dCas9-KRAB SUDHL-1 cells, qPCR confirmed significant depletion of guide RNA-targeted genes (SEL1L and DPH1), with greater than 16-fold depletion relative to cells expressing scrambled guide RNAs (**Supplementary Figure 4A**).

### Single guide RNA (sgRNA) lentiviral library transduction

hCRISPRi-V2 (RRID: Addgene_83969) is a commercially available guide RNA library that targets 18,905 protein coding genes with five independent guides targeting alternate promoter regions of each gene, and 1895 scrambled guide RNA controls that have no target. This library was a gift from Jonathan Weissman (http://n2t.net/addgene:83969). hCRISPRi-v2 was transfected into HEK293T cells for lentiviral packaging by the same protocol described above for dCas9.

### CRISPRi screens with PTCL culture under drug treatment

SUDHL-1-dCas9-KRAB cells were infected with lentivirus-packaged hCRISPRi-V2 guide RNA library at multiplicity (MOI) <0.4. This infection was performed by overnight incubation of 5 100mL replicate flasks each containing 750,000 live SU-DHL-1 dCas9-KRAB cells per mL in complete HPLM plus 0.8 μg/mL polybrene (Catalog number: Sigma TR-1003). Starting 48 hours after infection, 0.2 μg/mL puromycin was added to each flask for five days to select for infected cells. On day 5, flow cytometry confirmed that >90% of cells were infected according to significantly greater tagBFP signal. Cultures were transferred to complete HPLM without puromycin and grown for 24 hours prior to the addition of therapies for genome wide CRISPR screens.

Immediately prior to drug treatments, 120 million cells were harvested and frozen at –80 °C to serve as the time zero (T0) sample to assess guide RNAs initial frequencies. Cells were dispensed into two to three 175 cm^2^ flasks per treatment group, each with a density of 500,000 live cells per mL in 120mL of complete HPLM. CRISPR interference screens under different drugs were carried out in parallel in this scheme, with a common time zero sample serving as the baseline for multiple trug treatment groups. Alongside treatment groups, a DMSO-treated control flask was populated and harvested at the same final time point. All control flasks were consistently exposed to 0.1% DMSO by volume for the duration of the experiment. Cell growth was monitored daily using the average of multiple live cell counts using Trypan Blue and a Bio-Rad TC20 Automated Cell Counter (RRID: SCR_025462). Doses of each therapy were administered in pulses to each flask over 14 days to achieve a 2^8^-2^10^ fold depletion of cell growth compared with the DMSO control, with 40-60% of cells killed by each pulse drug administration (**Supplementary Figure 5**).

After treatment, 120 million cells were harvested from cultures in each treatment group. Genomic DNA was extracted from 80 million cells from each group (QIAGEN DNeasy blood and tissue kit). Library preparation was performed by amplifying guide RNAs contained in 400 μg of genomic DNA from each harvest. Amplification was performed in 60-84 parallel polymerase chain reactions using Q5 High Fidelity Master Mix (New England BioLabs Catalog number: m0492L) with primers that add Illumina adapters and individual indices to gRNAs from each treatment group. Amplified DNA representing the abundance of each guide RNA at T0 and at the end of each treatment period were pooled and sequenced with minimum 200x coverage by short read Illumina sequencing by the UNC High Throughput Sequencing Facility (HTSF, RRID: SCR_022620) for the romidepsin, pralatrexate, gemcitabine, and oxaliplatin screens and the Novogene Corporation in Sacramento, CA for the bortezomib screen. Phenotype scores and statistical significance were calculated using the analysis pipeline from *Horlbeck et al*^7^, based on the platform for phenotype scoring in pooled genetic screens from *Kampmann et al*^37^. Briefly, guide RNAs are assigned a ρ phenotype score that is the ratio of their abundance in the drug treated condition versus the DMSO-treated condition. When a ρ phenotype score is zero, that guide confers no change in sensitivity for a cell as compared to a wildtype cell. Alternatively, a guide with a score of 1 corresponds to complete resistance to drug treatment and negative guide scores are calculated for guides that sensitize cells to drug treatment relative to a wildtype cell. hCRISPRi-v2 has five replicate guide RNAs each targeting a different region of the promoter for a gene of interest for knockdown, and in the final comparison of guide RNAs across the library, the average phenotype score for the strongest three out of five guides per gene is measured to represent the effect of a gene of interest’s knockdown.

### Proteomic profiling of T– and NK-cell cultures

Total protein extraction was performed from 12 million cells of each lymphoma culture. Pelleted cells were triple rinsed in chilled, sterile PBS and then resuspended in cell lysis buffer. Cells were incubated on ice for 10 minutes and then probe-sonicated. Lysates were centrifuged and clear supernatant was collected. Bio-Rad microplate protein assay (Bio-Rad Catalog number: 500-0006) was used with a BSA standard curve to confirm total protein concentration for each lysate. 120 micrograms of total protein lysate for each culture were snap-frozen and submitted to the UNC Metabolics and Proteomics (MAP) core (RRID: SCR_001053).

Total protein samples were separated using a Thermo Vanquish Neo LC system (RRID: SCR_026495) and protein expression was quantified with an Orbitrap-Astral MS system (RRID: SCR_026205). For rigor and reproducibility, all protein samples were pooled and intermittently re-analyzed to test consistency in protein expression profiles for the pool across multiple runs (**Supplementary Figure 6**). In Spectronaut v19 proteins were aligned with the human protein database from Uniprot (Homo sapiens, Taxonomy ID: 9606). 10,682 proteins were identified, of which 87 were known common contaminants that were removed from analysis. Additional proteins were removed because they were ‘orphan peptides’ identified in only one lymphoma culture out of the panel, yielding roughly 9,200 proteins expressed consistently across the PTCL panel. A cross-run normalization was used to normalize protein abundance for each protein to the median expression for the set of all proteins. This median-normalized protein abundance was log-2 transformed for the association of protein expression with drug sensitivities in our analysis. We defined a minimum protein abundance threshold for a protein to be identified as detected, selecting the bottom 10^th^ percentile of abundance as non-detected. Proteins needed to be above this minimum abundance threshold in greater than 80% of the cell culture panel to be included in subsequent analysis, yielding 8181 total measured proteins for evaluation.

### Gene expression profiling of T– and NK-cell cultures

RNA extraction was performed from 1 million cultured cells for each T and NK culture using QIAGEN RNeasy Mini Kit (Catalog number: 74104) including on-column digestion of DNA with the QIAGEN RNase-Free DNase Set (Catalog number: 79254). cDNA Library Preparation and mRNA-sequencing were performed by the Novogene Corporation in Sacramento, CA. The top half of differentially expressed genes across the cell line panel were identified by rank-ordering gene expression for all transcripts by their standard deviation of expression. Correlations between gene expression (normalized counts) and drug sensitivity (AOC) were quantified by Spearman rank correlation.

### Candidate biomarker selection and sensitivity prediction

For each CRISPRi drug screen, genes were rank-ordered based on a combined score for their significance and phenotypic effect. This combined score is calculated as the absolute value of the average phenotype score for each gene multiplied by the negative Log of the Mann Whitney P-value that defines the difference between each gene’s distribution of gRNA phenotype scores and the distribution of phenotype scores measured for negative control guides. Further, after rank-ordering genes, the rankings of negative-control pseudogenes allowed us to infer a False Discovery Rate (FDR) for top ranked genes by calculating the ratio of pseudogene hits to real genes while progressing through the ranked list. All genes above the point in the ranked list where 10% of the hits were pseudogenes were considered to confer significant phenotype scores. Meaningful drivers of drug resistance and hypersensitivity were defined as those that were identified below the 10% FDR threshold for each CRISPR screen, up to a maximum Mann Whitney P-value of 0.05. These lists, of often greater than 100 CRISPRi-defined genes, were taken for each drug, and spearman rank correlations were calculated between the proteomic expression of these genes across diverse PTCL cultures. Only those genes that met independent criteria for correlation and causation were identified as candidate biomarkers.

### Leave-One-Out-Cross Validation to test prediction of drug sensitivity

We used a non-parametric approach to construct protein expression signatures for drug sensitivity prediction. For each cell culture, Z-scores of the selected protein features were summed to generate a composite score, giving equal weight to each feature, with a coefficient of +1 or –1 according to whether knockdown caused resistance or hypersensitivity. To evaluate predictive performance, we applied leave-one-out-cross validation (LOOCV). In each iteration of LOOCV, one culture was excluded, and a linear regression was performed between the summed Z-scores and measured drug sensitivities in the remaining cultures. The resulting regression was used to predict the drug sensitivity of the excluded culture from its sum of Z-scores. This process was repeated for each culture in the panel. Predictive accuracy was assessed by the Spearman rank correlation between observed and predicted drug sensitivities across all cell cultures.

### Simulated clinical trials of pralatrexate plus chemotherapy

Clinical trials of CHOP versus COP plus pralatrexate were simulated using a published ‘population tumor kinetics’ model, which simulates inter-patient and intra-tumor variation in response to multi-drug therapy and reproduces the clinically observed PFS of CHOP for Diffuse Large B-Cell Lymphoma^38^. The published model was re-calibrated to reproduce the PFS of CHOP for PTCL, observed in the Swedish Lymphoma Registry (n=755 patients)^39^ (**Supplementary Table 4**). A parameter describing average PTCL sensitivity to pralatrexate was fitted to reproduce the observed PFS of pralatrexate from pooled trials in relapsed/refractory PTCL^20^. Since the intended trial will replace doxorubicin (the H in CHOP) with pralatrexate due to overlapping toxicity^40^, we simulated COP plus pralatrexate, by setting doxorubicin dose to zero. As described in the prior publication^38^, biomarker-based patient selection was modelled using a parameter ‘rho’ that quantifies correlation between pralatrexate sensitivity and a biomarker, with rho = 0 denoting no predictive value, and rho = 1 denoting perfect selection of the most pralatrexate sensitive patients. We modeled rho = 0.6 to approximate the experimentally measured correlation between SLC19A1 protein expression and pralatrexate sensitivity in PTCL cultures.

### Statistical analyses

Statistical tests included spearman rank correlation of cell viabilities with the expression of biomarkers (**Figure 2**), Mann Whitney U-tests of cell viabilities between cultures predicted to be sensitive and insensitive to therapies (**Figure 3**, **Supplementary Figure 7**). Fitted sigmoidal dose response functions were computed in Mathematica 13 (**Figure 1**, **Supplementary Figure 2**, **Figure 3**, **Supplemental Figure 7**).

**Figure 1.**
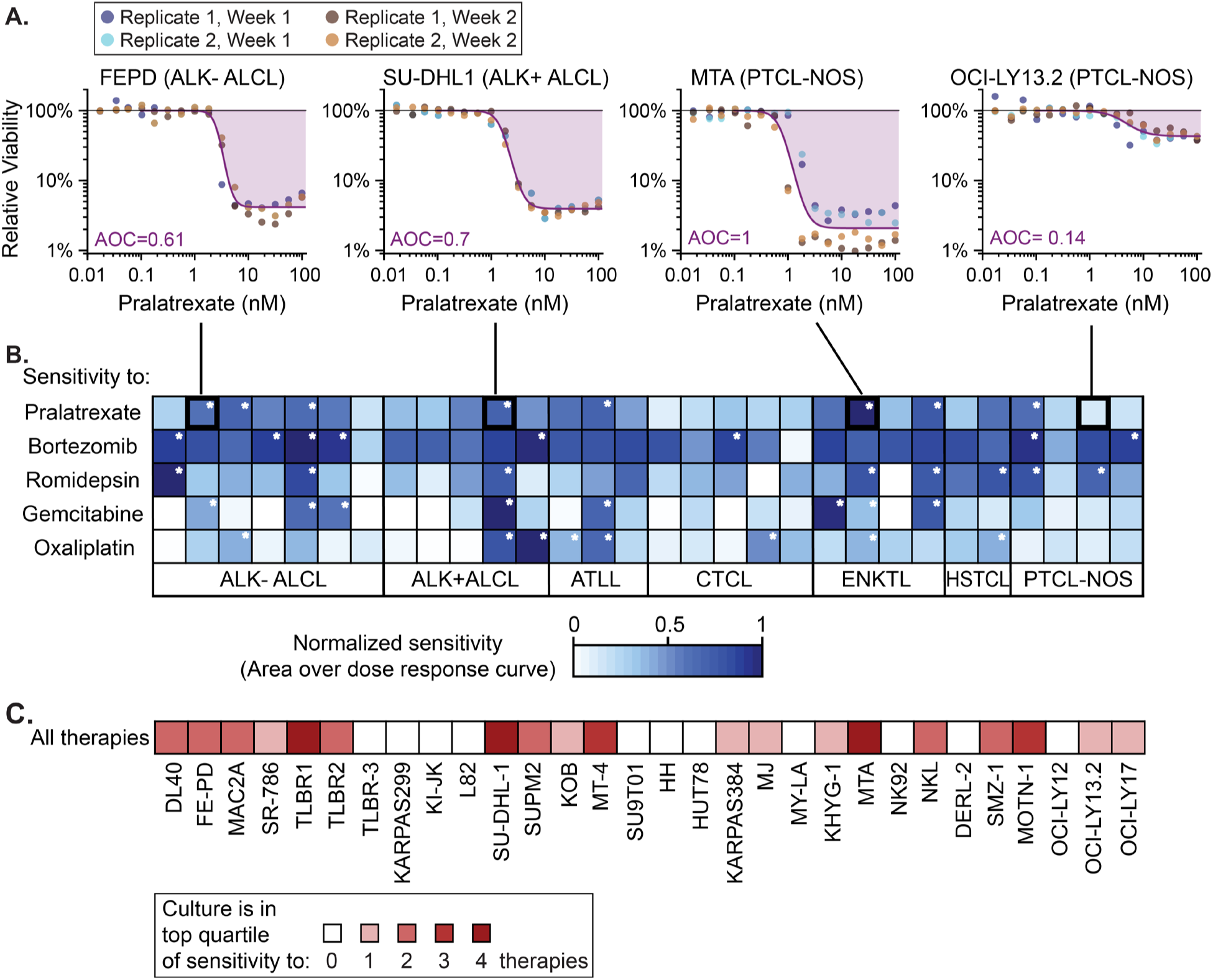
Most human T– and NK-cell lymphoma cultures are highly sensitive to at least one clinically active therapy. **A.** Example dose-response measurements of T– and NK-cell lymphoma cultures treated with pralatrexate for 72 hours. Cell viability was measured by luminescence (CellTiter-Glo) relative to untreated controls. Replicates included two independently propagated cultures, assayed on different plates across different weeks, yielding 3-4 replicates for each of 30 lymphoma cultures. **B.** Normalized drug sensitivities of 30 lymphoma cultures to 5 therapies used in the second-line treatment of PTCL. Sensitivities were quantified as the area over the dose-response curve (AOC), up to a clinically relevant maximum concentration (0.9 μM romidepsin^32^, 0.06 μM pralatrexate^33^, 0.8 μM bortezomib^34^, 0.1 μM gemcitabine^35^, and 4 μM oxaliplatin^36^). White asterisks mark cultures in the top quartile of sensitivity to each therapy. **C.** Heatmap indicating which lymphoma cultures are in the top quartile of sensitivity to one or more drugs.

**Figure 2.**
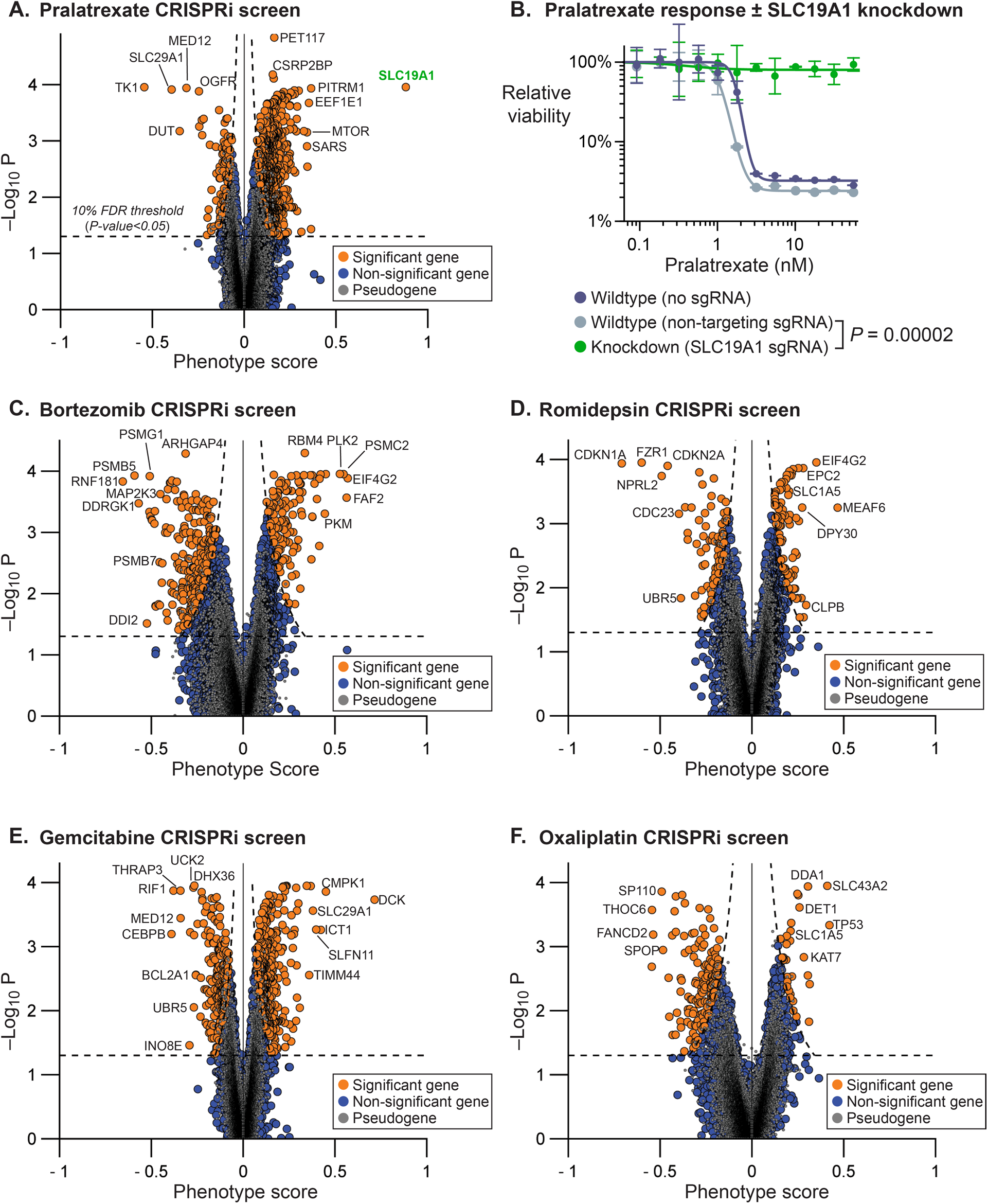
CRISPR interference screens identify genes whose knockdown alters sensitivity to second-line therapies for T-cell lymphomas. Volcano plots of phenotype scores and P values from genome-wide CRISPR interference screens under treatment with (**A**) pralatrexate, (**C**) bortezomib, (**D**) gemcitabine, (**E**) romidepsin, and (**F**) oxaliplatin. Orange points denote gene knockdowns with false discovery rate (FDR) below 10%, calculated from comparison with ‘pseudogene’ controls from non-targeting guide RNAs (gray; Methods); this threshold corresponds to Mann Whitney P values <0.05 (noting that FDR is not type 1 error). **B**. Effect of SLC19A1 knockdown on pralatrexate resistance. Pralatrexate dose response was measured in SU-DHL-1 cells expressing dCas9-KRAB, either untransformed, transformed with a non-targeting guide RNA, or transformed with a guide RNA targeting SLC19A1. Error bars show the standard deviation of four replicate measurements. At a physiologically relevant pralatrexate concentration (60nM, based on human pharmacokinetic studies) SLC19A1 knockdown abolished pralatrexate response compared with non-targeting control (*P* = 0.00002, Student’s t-test).

**Figure 3.**
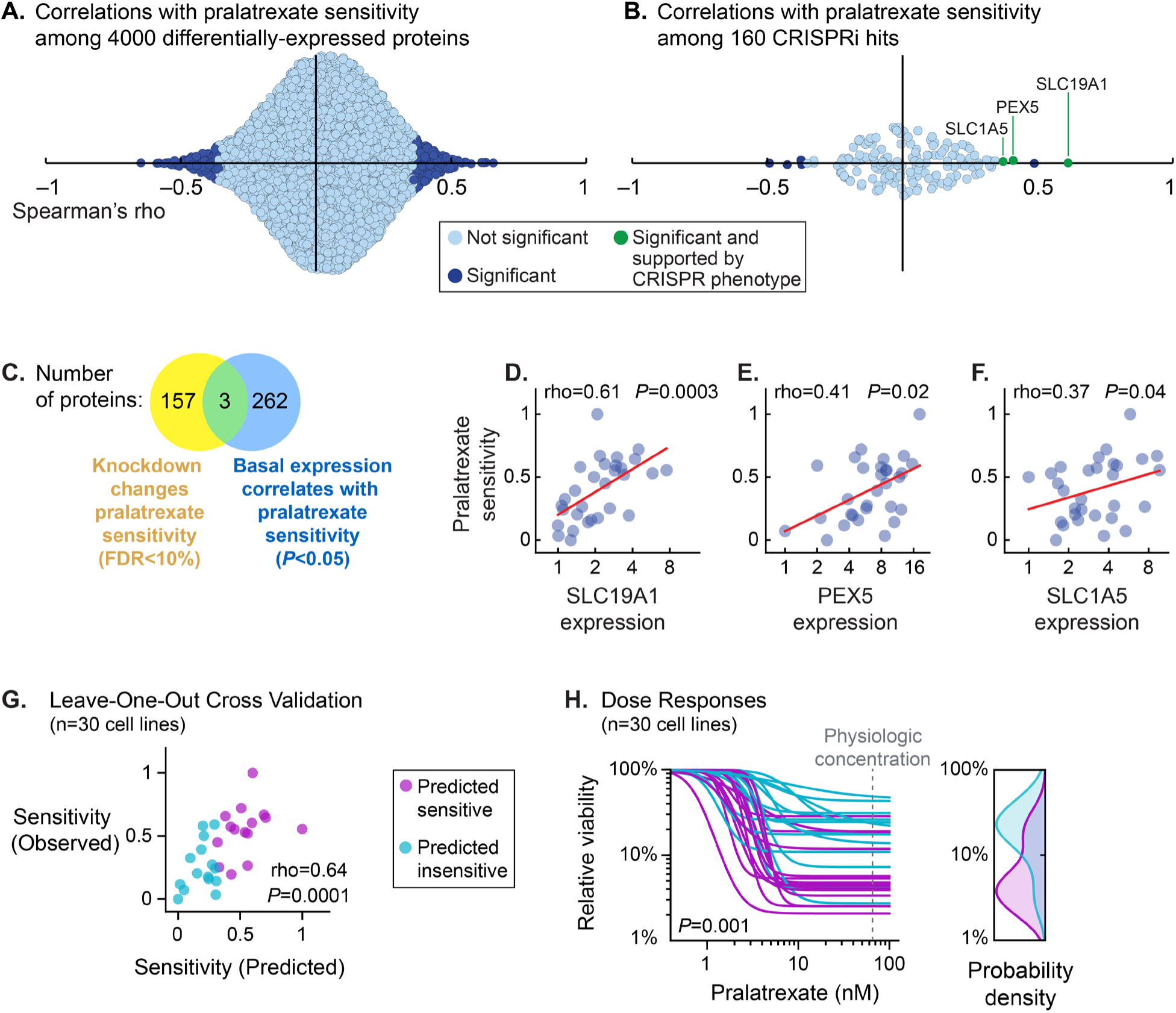
Integrating correlative associations and causal hits from CRISPR screens identifies features that predict drug sensitivities. **A.** Distribution of spearman rank correlations between pralatrexate sensitivity and expression levels of 4000 proteins that are differentially expressed across T– and NK-cell lymphoma cultures. Dark blue points mark hundreds of proteins with nominally significant associations (*P* < 0.05). **B**. Subset of the correlations in panel A, restricted to proteins whose corresponding genes were significant hits in the pralatrexate CRISPRi screen. Green points denote hits significant and concordant across both screens (e.g. significant positive correlation between protein expression and pralatrexate sensitivity, and knockdown confers significant resistance). **C**. Venn diagram depicting numbers of proteins whose knockdown significantly altered pralatrexate sensitivity in CRISPRi screens (yellow) and proteins whose abundance significantly correlated with pralatrexate sensitivity across PTCL cultures (blue). Three proteins in the intersection (green) meet both causal and correlative criteria. **D, E, F.** Relative expression of the 3 dual-criteria proteins (SLC19A1, PEX, SLC1A5) versus pralatrexate sensitivity (normalized AOC) across the 30 lymphoma cultures. Spearman’s rank correlation coefficients (rho) and P-values are shown. Red lines show linear regression. **G.** Scatterplot of observed pralatrexate sensitivities (AOC) versus predicted sensitivities from leave-one-out cross-validation of a protein expression signature (Methods). Predictive performance was quantified by Spearman’s rank correlation between observed and predicted AOC (rho=0.64, *P*=0.0001, n=30). Lymphoma cultures are colored according to predicted sensitivity above (purple) or below (cyan) the median. **H.** Dose-response curves for the 30 lymphoma cultures, stratified by predicted sensitivity from panel G. Cultures predicted to be more sensitive (purple) had significantly greater AOC than others (*P*=0.001, Mann Whitney U test, n=30), and exhibited greater potency and depth of inhibition; viability of predicted-sensitive cultures was significantly lower at a physiologically relevant pralatrexate concentration (*P*=0.02, Mann Whitney U test, n=30).

### Data availability

T– and NK-cell lymphoma culture drug sensitivities and protein and mRNA expression measurements are in **Supplementary File 1**. Data from CRISPR interference screens are available in **Supplementary File 2**.

## RESULTS

### PTCL Cells are differentially sensitive to second-line therapies

A number of therapies can elicit 20 to 30% response rates in relapsed/refractory PTCL. The potential utility of biomarker-guided treatment depends on whether different patients respond best to different therapies, or if some subset of patients is responsive to nearly all treatment options, with remaining patients resistant to all therapies. We therefore first asked whether different T– and NK-cell lymphoma cultures respond best to different drugs. Drug sensitivities were profiled in 30 lymphoma cultures using 16-point dose–response assays for each of five therapies: romidepsin, pralatrexate, bortezomib, gemcitabine, and oxaliplatin (**Figure 1A**). Drug responses measured by a luminescent viability assay exhibited a large dynamic range (from 100% to <0.1% relative viability) and were reproducible across replicate experiments on different weeks (average R^2^=0.85; **Supplementary Figure 1**), supported by stringent control of seeding densities and growth conditions (Methods).

Drug sensitivities, represented by fitted dose response curves (**Supplementary Figure 2A**), varied substantially between lymphoma cultures, with most being resistant to some therapies but sensitive to others. Nineteen of thirty (63%) cultures were in the top-quartile of sensitivity to at least one drug studied (**Figures 1B, 1C**), indicating a scenario where matching drugs to individual lymphomas could outperform a uniform treatment strategy. However, these differences were not clearly delineated by immunophenotype (**Supplementary Figure 2B-F**), and growth rates of cultures also did not correlate with sensitivity to any therapy (**Supplementary Figure 3**), supporting the idea that further molecular insights are required to predict drug sensitivity.

### Determinants of drug sensitivity are identifiable with genome-wide CRISPR interference screens

To identify genes whose expression level can directly modify drug sensitivity, we performed genome-wide CRISPR interference screens in SU-DHL-1 cells under treatment with either romidepsin, pralatrexate, bortezomib, gemcitabine, oxaliplatin, or DMSO as a no treatment condition. SU-DHL-1 was used because of its initial sensitivity to each drug of interest, and because it could be efficiently transduced by lentivirus; these features are essential for CRISPR screens for modifiers of drug sensitivity. Measurement of expression in representative target genes SEL1L and DPH1 by qPCR demonstrated greater than 16-fold gene knockdown (94% expression depletion) when guide RNAs against these genes were transduced into SU-DHL-1 cells expressing dCas9-KRAB (**Supplementary Figure 4A**). SU-DHL-1 cells expressing dCas9-KRAB were transduced with hCRISPRi-V2, a library of guide RNAs targeting 18,905 protein-coding genes with 5 guides per gene, and 1,895 non-targeting control guides (Addgene_83969). After transduction, cultures were propagated for 13 to 14 days under drug treatment schedules that inhibited proliferation by 8 to 10 doublings relative to DMSO-treated controls (**Supplementary Figure 4B-F**), which in principle allows a completely resistant clone to be enriched by a factor of 2^8^ to 2^10^. Cells were thereafter harvested and abundance of clones bearing each guide RNA were counted by next-generation sequencing (Methods).

For each drug, hundreds of gene knockdowns conferred significant resistance or hypersensitivity. To account for multiple hypothesis testing, hits were defined by False Discovery Rate (FDR) < 10% in comparison to non-targeting controls, corresponding to nominal *P*-value cutoffs < 0.029 to 0.049, as described^41^ (**Figure 2A, C-E**, **Supplementary Table 2**). In an exploratory analysis of resistance mechanisms, STRING analysis of significant CRISPR hits highlighted relationships such as knockdown of proteasome subunits (PSMB1-4, PSMC1-5) conferring resistance to bortezomib, and knockdown of genes in the Fanconi anemia DNA repair pathway enhancing sensitivity to oxaliplatin^42^ (**Supplementary Table 2**). Additionally, several genes previously identified as drivers of drug resistance in other cancers were reproduced in T-cell lymphoma, including SLC43A2 knockdown conferring resistance to oxaliplatin^43^ and BTG1 and NFE2L1 knockdown enhancing sensitivity to bortezomib^44^. The strongest resistance phenotypes arose from knockdown of the reduced folate carrier SLC19A1 for pralatrexate and deoxycytidine kinase (DCK) for gemcitabine, consistent with known mechanisms: SLC19A1 mediates pralatrexate uptake into cells^30,45^, and DCK is required to phosphorylate gemcitabine for its incorporation into DNA and induction of DNA damage^46^.

Single-gene knockdown experiments validated phenotypes of top-scoring genes from the multiplexed CRISPRi screen. SU-DHL-1 cells expressing dCas9-KRAB were transduced with single-guide RNAs targeting individual genes, or non-targeting controls, before conducting dose–response assays. Knockdown of SLC19A1 conferred complete resistance to pralatrexate at concentrations 100 times greater than the wildtype IC50 (**Figure 2B**). Knockdown of SLFN11 conferred resistance to gemcitabine (**Supplementary Figure 5A**), and it is well-established in prior literature that SLFN11 is a critical determinant of activity of DNA damaging agents^47^. Knockdown of CDKN1A heightened sensitivity to histone deacetylase inhibitors romidepsin and belinostat, consistent with prior work in B-cell lymphomas^48^ (**Supplementary Figure 5B-C**). These single-gene perturbations confirm the ability of a multiplexed screen to identify genes whose expression influences drug sensitivity, providing a short-list of candidates for correlative analyses across a panel of PTCL cultures.

### Proteomic profiling of PTCL Cultures identifies features significantly correlated with drug sensitivity

We next measured basal protein expression for the panel of 30 T– and NK-cultures to identify proteins whose abundance significantly correlate with drug sensitivities, as measured with dose response experiments in Figure 1. Bulk protein expression profiles were collected from each lymphoma culture by LC/MS-MS. Only proteins quantifiable in at least 80% of cultures were included in subsequent analysis, yielding 7,920 proteins (Methods). Protein abundance measurements were strongly consistent across independently propagated flasks of the same line (R^2^=0.98; **Supplementary Figure 6A**). Generally the proteomes of these cell lines were correlated, with a minimum spearman correlation of 0.8, indicating an expected level of similarity due to these lymphomas being derived from similar tissues of origin (**Supplementary Figure 6B**).

To identify proteins whose expression correlates with drug sensitivity, only the top half of differentially expressed proteins were considered (4000 total, according to the standard deviation of log_2_(relative abundance)), reasoning that homogeneously expressed genes are unlikely to explain heterogeneity in drug response. To illustrate the challenge posed by many candidate features, we first show that hundreds of proteins exhibit nominally significant correlations with drug sensitivity across the culture panel (**Figure 3A**, **Supplementary Figure 7A-D**). This motivates the combination of these data with CRISPR interference screens to identify a restricted set of functionally influential candidates.

To focus biomarker discovery on causally relevant candidates, we next assessed correlations only amongst genes whose knockdown by CRISPRi significantly perturbed drug response, and whose protein product was differentially expressed across cultures. Features could influence drug sensitivity in either direction: knockdown could cause resistance and protein abundance positively correlates with sensitivity, or knockdown could cause hypersensitivity and protein abundance negatively correlates with sensitivity. Whereas a genome-wide search found hundreds of hits per drug, this focused search identified 2 to 5 genes per drug that significantly correlate with drug response across the panel of PTCL cultures, representing the intersection of causal and correlative search criteria (**Figure 3B-F** for pralatrexate; **Supplementary Table 3 and Supplementary Figure 7A-D** for other therapies).

### Protein expression signatures predict sensitivity of T– and NK-cell lymphoma cultures to 5 clinically active therapies

Since multiple genes satisfied the above criteria for each drug, we tested whether multi-gene expression signatures could distinguish sensitive from resistant cultures. For each drug we created a simple non-parametric score by summing the Z-scores of candidate proteins, which represent how much a protein’s abundance in one culture deviates from its average across all cultures, with positive or negative sign according to whether knockdown caused resistance or hypersensitivity.

We tested how well protein expression signatures associate with drug responses using Leave-One-Out-Cross Validation (LOOCV). First, each culture was iteratively excluded from the dataset. A linear regression was fitted to the sums of Z-scores and observed drug sensitivities (Area Over the dose response Curve; AOC) for the remaining 29 cultures. This regression was used to predict the sensitivity of the left-out culture. For all five drugs, protein expression signatures yielded significant rank correlations between predicted and observed drug sensitivities for the set of cultures. The strongest prediction was made for pralatrexate (Spearman’s rho = 0.64, *P* = 0.0001; **Figure 3G**). Finally, cell lines were stratified into two groups according to greater than or less than median score, and in each case the higher scoring group of cultures was significantly more drug sensitive (Mann Whitney *P* values ≤ 0.03; **Figure 3H**, **Supplementary Figure 7E-F**).

### Transcriptomic profiling of PTCL demonstrates the utility of RNA level as a proxy for drug sensitivity prediction

Following the identification of candidate biomarkers by proteomics, the mRNA expression of the same genes was assessed for their potential use in drug sensitivity prediction. This assessment was performed to establish the potential for RNA level to serve as a proxy in cases where protein abundance for a biomarker of interest cannot be readily measured or exhibits insufficient dynamic range by, for example, immunohistochemistry. Drug sensitivity prediction by RNA expression was less accurate than by proteomics, though leave-one-out cross-validation using the same gene lists showed significant prediction of sensitivity to pralatrexate (r=0.61, *P*=0.0003), gemcitabine (r=0.50; *P*=0.005), and bortezomib (r=0.47, *P*=0.01), but not oxaliplatin or romidepsin (**Supplementary Figure 8**). Notable features including SLC19A1 are correlated with pralatrexate sensitivity by transcript expression as well as by protein level, but the correlation is less strong, indicating that SLC19A1 protein level may be a more robust predictor of pralatrexate efficacy.

### Simulating the potential for pralatrexate to improve patient outcomes in combination with CHOP

We simulated clinical trials to investigate whether patient selection with a predictive biomarker could increase the chance that pralatrexate improves Progression-Free Survival (PFS) when added to first-line combination therapy for PTCL. Because SLC19A1 mediates pralatrexate entry into lymphoma cells, variation in its expression provides a biologically plausible basis for biomarker-guided enrollment (**Figure 4A**), prompting us to simulate whether a biomarker as accurate as SLC19A1 could improve trial outcomes. Clinical trials were simulated by adapting a published model of the clinical efficacy of CHOP and RCHOP for DLBCL, which was previously shown to reproduce the PFS hazard ratios of various ‘RCHOP+X’ trials based on the monotherapy efficacies of the added agents in relapsed/refractory disease^38^. The model was calibrated to reproduce the observed PFS distribution of CHOP for previously untreated PTCL (**Figure 4B**)^39^, and the observed PFS distribution of pralatrexate monotherapy for relapsed/refractory PTCL (**Figure 4C**)^20^. This calibrated model was then applied to simulate an upcoming trial comparing CHOP versus COP plus pralatrexate, in which doxorubicin (H) is omitted from the investigational arm due to overlapping toxicity with pralatrexate^40^.

**Figure 4.**
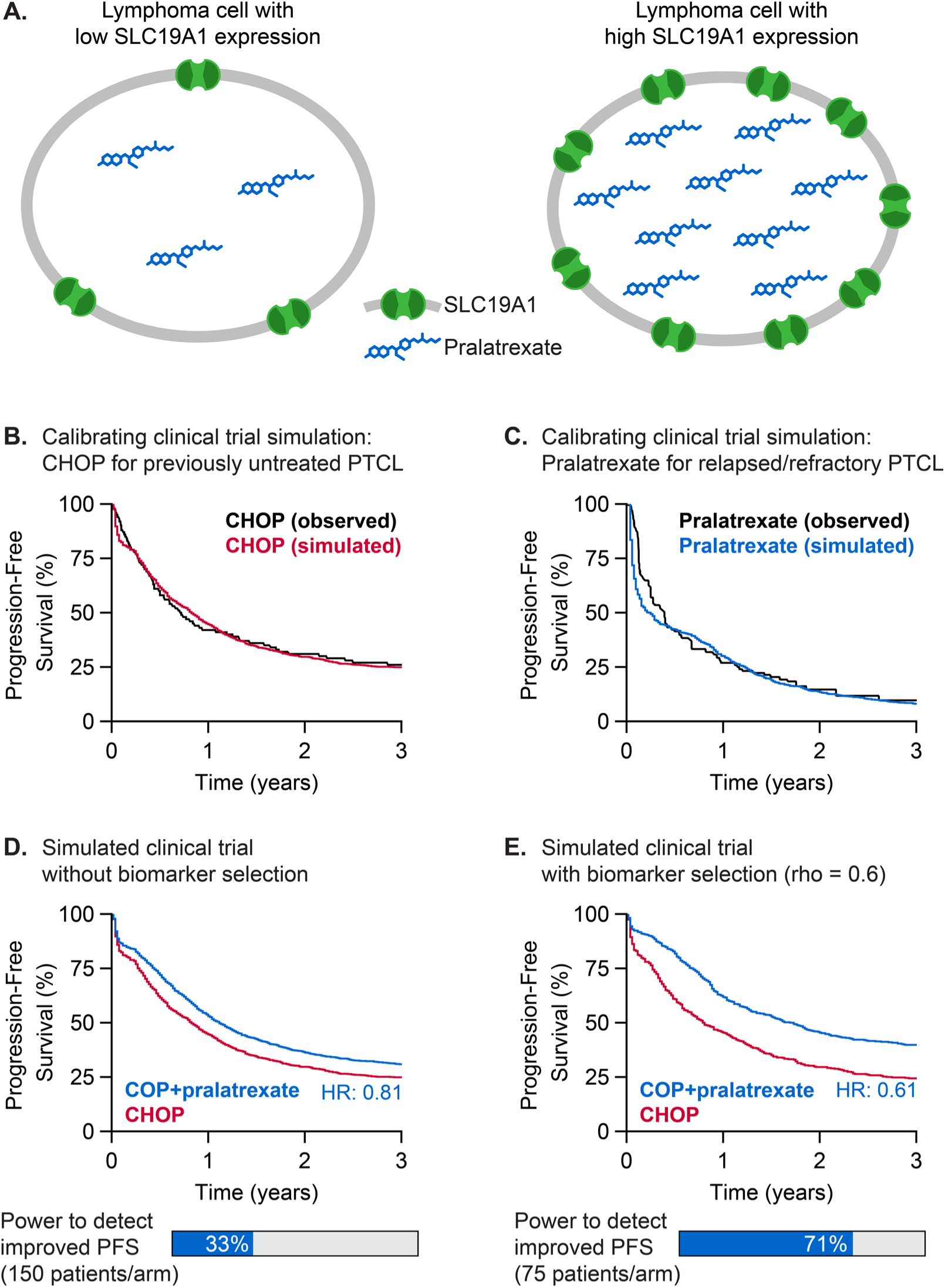
Simulated trials predict that biomarker-based patient selection increases the likelihood that pralatrexate improves first-line outcomes. **A.** Proposed mechanism by which SLC19A1 expression predicts pralatrexate sensitivity: lymphoma cells with lower or higher SLC19A1 expression accumulate lesser or greater intracellular pralatrexate concentrations, respectively. **B.** A published population-tumor kinetics model was calibrated to reproduce Progression Free Survival (PFS) outcomes for previously untreated PTCL treated with CHOP (Swedish Lymphoma Registry; Methods). **C.** The model was also calibrated to reproduce PFS of pralatrexate monotherapy for relapsed/refractory PTCL (pooled analysis of pralatrexate studies; Methods). **D.** Fitted parameters for pralatrexate and CHOP were applied to simulate COP+pralatrexate vs CHOP for previously untreated PTCL. Simulated trials without biomarker selection predicted PFS hazard ratio 0.81, and 33% power to detect significant benefit (α=0.05) with 150 patients/arm. **E**. Simulated trials employing biomarker-based patient selection, using a biomarker as correlated with pralatrexate sensitivity as SLC19A1 expression (rho = 0.6), predict improved PFS from COP+pralatrexate (hazard ratio 0.61) and 71% power to detect significant benefit with 75 patients/arm.

We first simulated trials with 1,000 patients per arm to estimate treatment effect with minimal sampling error (noise) and next subsampled these populations to assess how realistic sample sizes affect the statistical power to detect benefit. The predicted PFS hazard ratio for pralatrexate-COP versus CHOP was 0.81 (**Figure 4D**). However, in simulated trials with 150 patients per arm without biomarker selection, consistent with the intended design, only 33% of trials exhibited significantly improved PFS, predicting low statistical power to detect benefit in an unselected population. Conversely, simulated trials that enroll a subset of patients most likely to be sensitive to pralatrexate, according to a biomarker with predictive accuracy equal to that exhibited by SLC19A1 in PTCL cultures (rho=0.6), predicted a PFS hazard ratio of 0.61 (**Figure 4E**). In this case, trials of 75 patients per arm (reflecting a biomarker-selected subset) achieved significance in 71% of simulations, a much improved statistical power. Importantly, this simulation did not hypothesize a perfect biomarker: merely enriching for pralatrexate-sensitive lymphomas doubled statistical power with half as many patients. These simulations support the value of biomarker-guided enrollment to improve the chance of success of novel drug combinations.

## DISCUSSION

Biomarker-guided cancer treatment can improve outcomes by matching patients with therapies more likely to be effective for them. Correlative pharmacogenomic analyses that link molecular features (e.g. expression, mutations) to drug sensitivity yield important insights but often return large sets of associations due to testing vast numbers of features. Functional genomic screens provide complementary, causal evidence by identifying genes whose perturbation directly alters drug response. However, hits from such screens cannot all be expected to be useful biomarkers because their effect may be context-dependent, and they may not have relevant variation among patients. In this study, integrated correlative and functional genomic approaches overcome these limitations and identify manageable lists of candidate biomarkers for therapies used in the treatment of PTCL.

The reduced folate carrier SLC19A1 emerged as the strongest predictor of pralatrexate sensitivity in PTCL. Pralatrexate was designed to have higher affinity to SLC19A1 than methotrexate, so its mechanistic role in antifolate pharmacology is well established^49^. However, SLC19A1 has not been investigated as a biomarker to select patients for pralatrexate treatment. Previous studies of anti-folate chemotherapy identified loss of SLC19A1 as a mechanism of acquired resistance^50,51^ and germline SLC19A1 polymorphisms as predictors of methotrexate toxicity to normal tissues^52^. Prior studies of pralatrexate in various solid tumor cultures suggested that SLC19A1, folyl-polyglutamate synthase (FPGS), and dihydrofolate reductase (DHFR) expression might all serve as biomarkers of pralatrexate efficacy^51^. In the context of PTCL, only SLC19A1 showed predictive value, and its phenotype upon knockdown far surpassed the phenotypes of these other features (rho = 0.88 for SLC19A1; rho = 0.15 for FPGS; rho = –0.03 for DHFR). Thus, among mechanistically-linked features, only SLC19A1 satisfies orthogonal criteria for causation and correlation, and therefore its basal expression might provide the means to predict pralatrexate sensitivity in PTCL.

SLC19A1’s strong performance as a predictor highlights the broader influence of membrane transporters upon cancer cells’ sensitivity to small molecules. Solute carrier (SLC) and ATP-binding cassette (ABC) transporters are key determinants of cellular uptake and efflux, especially for drugs that structurally resemble endogenous metabolites^53,54^. Our CRISPR screens identified several transporters with the potential to predict drug sensitivities in T-cell lymphoma models.

Along with SLC19A1 for pralatrexate, SLC2A3 abundance was a candidate biomarker of gemcitabine sensitivity in PTCL cultures. SLC2A3, also known as glucose transporter 3 (GLUT3), mediates the uptake of multiple hexoses as well as the nucleoside analog capecitabine which contains a ribose moiety. Gemcitabine similarly contains a modified deoxyribose, and SLC2A3 upregulation was recently observed to drive gemcitabine sensitivity in pancreatic cancer cells^55^, suggesting SLC2A3 as a mediator of gemcitabine uptake. Additionally, SLC29A1, also known as the Equilibrative Nucleoside Transporter Member 1 (ENT1), is a known importer of gemcitabine with prognostic significance in gemcitabine-treated pancreatic cancer^56,57^, and its knockdown in PTCL brought about significant gemcitabine resistance (rho=0.4, *P* = 0.0003). These findings, together with the established roles of ABC transporters on chemosensitivity^58^, highlight importers and exporters as powerful determinants of small molecule sensitivity in cancers. Membrane transporters therefore warrant particular attention in biomarker discovery for small molecule therapies, and focused CRISPR libraries to perturb SLC and ABC families offer a tool to do so^59^.

The focus of this study was combining experimental methods to identify features for drug response biomarkers. Our approach to response prediction using the identified features was intentionally simple and parameter-free. More advanced computational approaches might improve response prediction, especially considering advances in machine learning. To support the application of such approaches to PTCL, we provide genome-wide CRISPR interference phenotypes, together with drug responses, transcriptomic, and proteomic profiles of the panel of PTCL cultures (**Supplementary Files 1 and 2**).

In this study we profiled drug sensitivities and protein expression for nearly all commercially available cell culture models of T– and NK-cell lymphoma. Thirty cultures is a modest sample size considering the extensive heterogeneity of T-cell lymphomas, emphasizing the need for validation in patient cohorts. A chief use of such preclinical work is to generate a biomarker hypothesis that might be confirmed in patient data, in contrast to using the same patient data for a ‘hypothesis generating’ analysis which would then require a second independent patient cohort for confirmation. Although drugs studied here are clinically active, preclinical activity or biomarkers thereof may not translate to clinical activity due to limitations in faithfully modeling physiological conditions *in vitro*. For example, supra-physiological folate levels in culture media may heighten dependence on DHFR^60^. However, recent trials in hematological malignancies have found that drug responses of *ex vivo* samples can translate to clinical responses. The EXALT trial showed that tailoring therapy based on *ex vivo* drug response profiles improved Progression-Free Survival compared to the prior line of therapy for 54% of patients with relapsed or refractory leukemias or lymphomas^61^. The ‘SMARTrial’ similarly generated *ex vivo* drug response profiles, and reported a case of a patient whose lymphoma cells resisted nearly every drug tested except pralatrexate, and the patient then responded to treatment by another DHFR inhibitor, methotrexate^62^. These observations support the translational relevance of measuring the sensitivities of lymphoma cells to drugs with known clinical efficacy against lymphoma. Some cancer therapies are active in patients but not in culture; our approach would not apply to such drugs, though focused CRISPR screens are feasible in mice. Not all candidate biomarkers identified here have clear mechanistic links to drug action. Although supported by knockdown phenotypes and correlative evidence, such hits may be either novel biology or false discoveries, and further characterization is warranted. Overall our preclinical studies demonstrate how functional genetic screens can narrow the hypotheses tested in correlative studies, and in principle, *in vitro* functional screens (or similar screens in mice) could also assist correlative analyses of clinical datasets.

Discovering biomarkers of drug response remains challenging, but combining functional genetic screens with correlative analyses can make the problem more tractable by narrowing the search space and prioritizing candidates supported by causal evidence. The inherent challenge of correlative analyses was demonstrated by the JAVELIN Renal 101 trial of tyrosine kinase inhibition with or without PD-L1 inhibition, where analysis of thousands of gene-survival associations produced hundreds of hits, but inevitably, none remained significant after correcting for multiple testing^63^. Functional screens highlight the subset of genes whose function directly influences drug response, and when integrated with correlative data across diverse models or patient samples, can enrich for biomarkers with true predictive value. Finally, simulated clinical trials confirm for our specific case what is broadly known to be true: for a therapy like pralatrexate whose efficacy is evidently limited to a subset of patients, employing biomarker-guided patient selection could substantially increase the likelihood of improving clinical outcomes.

## Authors’ Contributions

J.C.P: Conceptualization, investigation, formal analysis, writing. A.E.P: Trial simulations, writing. A.C.P: Conceptualization, formal analysis, writing.

## Conflict of Interest Disclosures

A.C.P. has received consulting fees from AstraZeneca, Kymera, Merck, Novartis, and Sanofi, research funding from Merck and Prelude Therapeutics, and equity in Oncko. A.E.P. has received consulting fees from Respiratorius AB.

## Funding

This work was supported by NIH grant R01CA279968.

## Acknowledgements

We thank David Weinstock for T– and NK-cell lymphoma cultures, the UNC Flow Cytometry Core (Ayrianna Hedgecock, Janet Dow, and Roman Bandy) for cell sorting, Novogene (Soumi Joseph) for assistance with CRISPRi library sequencing, and the UNC Michael Hooker Metabolomics and Proteomics core for LC/MS-MS (Laura Herring, Thomas Webb, Aurora Caberea, Natalie Barker-Krantz). This work was supported by NIH grant R01CA279968.

