## Supplementary Figures for "Integrating functional genomics and proteomics identifies Folate Carrier SLC19A1 as a predictor of pralatrexate sensitivity in T-cell lymphoma"

**Supplement Contents**


i. **Supplementary Table 1. T- and NK-cell lymphoma cultures and their sources**

ii. **Supplementary Figure 1. Reproducibility of dose response measurements in 30 T and NK Lymphoma Cultures**

iii. **Supplementary Figure 2. Immunophenotype-defined subgroups do not distinguish drug sensitivities for Mature T and NK cultures**

iv. **Supplementary Figure 3. TCL Culture Growth Rate has no significant correlation with sensitivity to second line therapies**

v. **Supplementary Figure 4. CRISPR Interference knockdown validation and growth projections for CRISPR Interference Screens with SU-DHL-1**

vi. **Supplementary Table 2. Summary of Gene Knockdowns with significant phenotype scores in CRISPRi screens**

vii. **Supplementary Figure 5. Validation of CRISPR interference knockdowns' impact on drug sensitivity of ALK+ ALCL culture**

viii. **Supplementary Figure 6. Proteomic profiles of 30 T and NK lymphoma cultures show a high degree of similarity**

ix. **Supplementary Figure 7. Identification of candidate biomarkers for additional second line therapies beyond pralatrexate**

x. **Supplementary Table 3. Candidate biomarkers of drug sensitivity to second line therapies and literature support for their mechanistic relationships**

xi. **Supplementary Figure 8. mRNA Expression of candidate biomarkers is similarly predictive of drug sensitivity in some cases**

xii. **Supplementary Table 4. Parameter values for clinical trial simulations**

xiii. **Supplementary References**

**Supplementary Table 1. T- and NK-cell lymphoma cultures and their sources**

| Cell Line | Source and RRID | Histology-Defined Subtype | Clinical Information for Patient Source | STR Profile Match (%) | Culturing Conditions |
|---|---|---|---|---|---|
| DERL-2 | DSMZ (CVCL_2016) | HSTCL | 30 y.o. male[1] | 100 | Requires Recombinant IL-2 |
| DL40 | JCRB (CVCL_2889) | ALK- ALCL | 64 y.o. female[2] | 100 | Standard |
| FE-PD | Weinstock (CVCL_H614) | ALK- ALCL | 45 y.o. female[3] | NA | Standard |
| HH | ATCC (CVCL_1280) | CTCL | 61 y.o. male[4] | 98 | Standard |
| HUT78 | ATCC (CVCL_0337) | CTCL | 58 y.o. male[5] | 100 | Standard |
| Karpas299 | Sigma (CVCL_1324) | ALK+ ALCL | 25 y.o. male[6] | 100 | Standard |
| Karpas384 | Sigma (CVCL_2541) | CTCL | 48 y.o. male[7] | NA | Standard |
| KHYG-1 | JCRB (CVCL_2976) | ENKTL | 45 y.o. female[8] | 98.0 | Requires Recombinant IL-2 |
| KIJK | DSMZ (CVCL_2093) | ALK+ ALCL | 15 y.o. male[9] | 100 | Standard |
| KOB | Weinstock (CVCL_VN49) | ATLL | 72 y.o. male[10] | NA | Requires Recombinant IL-2 |
| L82 | DSMZ (CVCL_2098) | ALK+ ALCL | 24 y.o. female[11] | 96.4 | Standard |
| MAC2A | Weinstock (CVCL_H637) | ALK- ALCL | 31 y.o. male[12] | NA | Standard |
| MJ | ATCC (CVCL_1414) | CTCL | 50 y.o. male[13] | 98 | Standard |
| MOTN-1 | DSMZ (CVCL_2127) | PTCL-NOS | 63 y.o. female[14] | 100 | Requires Recombinant IL-2 |
| MT-4 | DSMZ (CVCL_2632) | ATLL | 50 y.o. male[15] | 97.9 | Standard |

| | | | | | |
|---|---|---|---|---|---|
| MTA | JCRB (CVCL_3032) | ENKTL | 58 y.o. female[16] | 96.8 | Requires 20% FBS |
| MY-LA | Sigma (CVCL_M415) | CTCL | 82 y.o. male[17] | 92.8 | Requires Recombinant IL-2 and IL-4 |
| NK-92 | DSMZ (CVCL_2142) | ENKTL | 50 y.o. male[18] | 100 | Requires 20% FBS and Recombinant IL-2 |
| NKL | Weinstock (CVCL_0466) | ENKTL | 62 y.o. male[8] | NA | Requires Recombinant IL-2 |
| OCILY12 | UHN (CVCL_8796) | PTCL-NOS | 37 y.o. male[19] | NA | Standard |
| OCILY13.2 | UHN (CVCL_8797) | PTCL-NOS | 28 y.o female[19] | NA | Standard |
| OCILY17 | UHN (CVCL_8798) | PTCL-NOS | 72 y.o. male[20] | NA | Standard |
| SMZ-1 | DSMZ (CVCL_RL84) | HSTCL | 46 y.o. male[21] | NA | Standard |
| SR-786 | DSMZ (CVCL_1711) | ALK- ALCL | 11 y.o. male[22] | 100 | Standard |
| SU9T01 | Weinstock (CVCL_B7P5) | ATLL | no pt information[23] | NA | Requires Recombinant IL-2 |
| SU-DHL-1 | DSMZ (CVCL_0538) | ALK+ ALCL | 10 y.o. male[24] | 100 | Standard |
| SUPM2 | DSMZ (CVCL_2209) | ALK+ ALCL | 5 y.o. female[25] | 100 | Standard |
| TLBR-1 | DSMZ (CVCL_L177) | ALK- ALCL | 42 y.o. female[26] | 98.3 | Requires Recombinant IL-2 |
| TLBR-2 | DSMZ (CVCL_A1EY) | ALK- ALCL | 43 y.o. female[27] | 100 | Standard |
| TLBR-3 | DSMZ (CVCL_A1EZ) | ALK- ALCL | 45 y.o. female[27] | 98.1 | Requires Recombinant IL-2 |

DSMZ=Deutsche Sammlung von Mikroorganismen, Leibniz Institute; JCRB=Japanese Collection of Research Bioresources; ATCC=American Type Culture Collection; UHN=University Health Network, Toronto Canada[19,20]; “Weinstock”= originally obtained by the lab of David Weinstock[28]; An “NA” in the STR profile match field indicates that the culture is not listed in the Cellosaurus database of STR profiles.

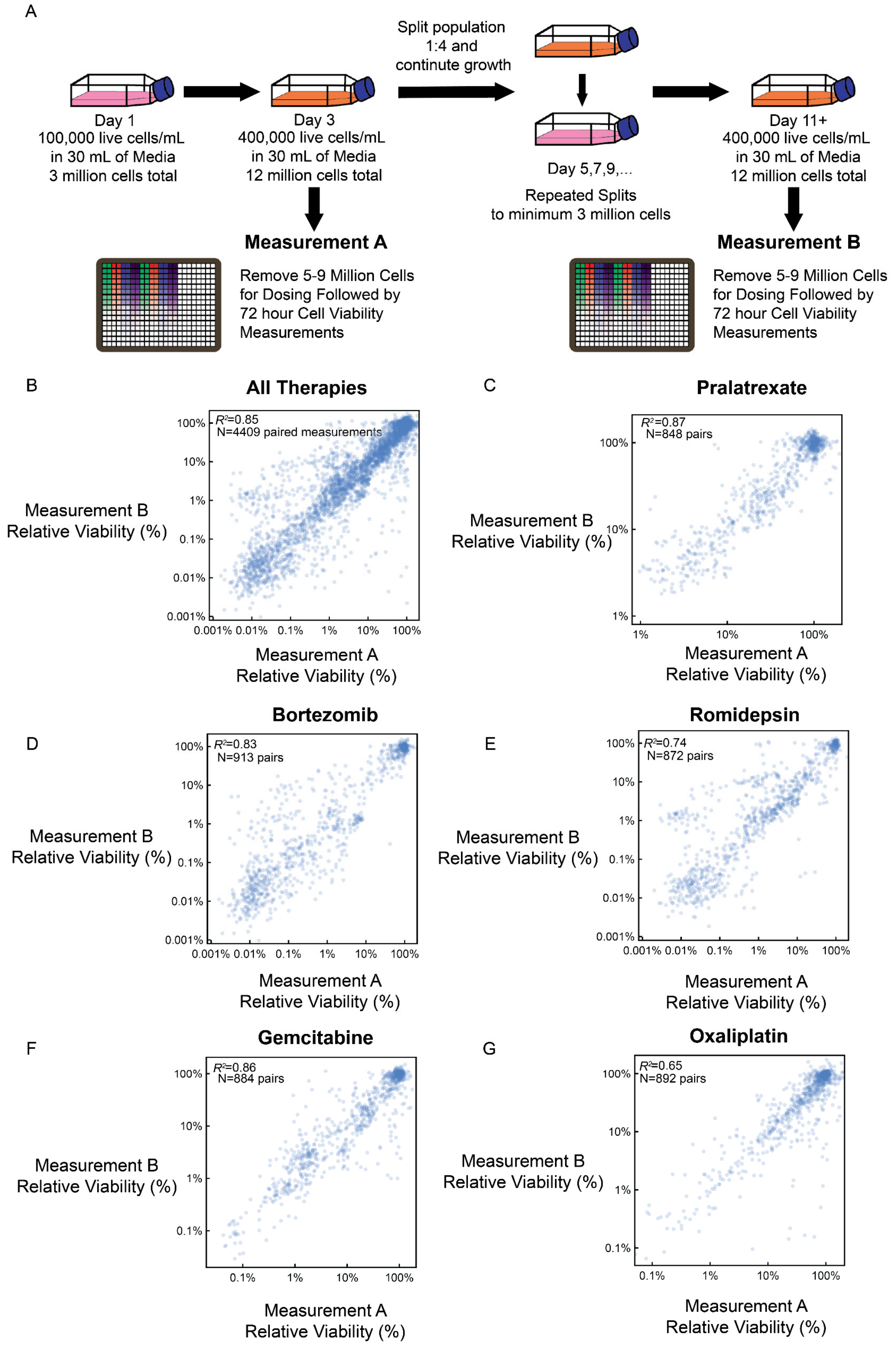


**Supplementary Figure 1. Reproducibility of dose response measurements in 30 T and NK Lymphoma Cultures.** Panel A contains a schematic of the process used to collect dose response measurements in biologic replicates across T and NK lymphoma cell cultures. Cultures were regularly passaged in 30mL of culture to a minimum density of 100,000 live cells /mL of media prior to the collection of dose responses. The schematic's representation of a consistent 1:4 split from 400,000 live cells per mL to 100,000 live cells per mL serves as a common example of culture treatment, considering the majority of T and NK cell cultures exhibit an approximate 24 hour doubling time, but the actual dilution of cultures at each time point was dependent upon measured live cell density immediately prior to passaging. Dose responses were collected in 384-well plates at two points, Measurement A and Measurement B, with a minimum of one week of routine passaging to evaluate the reproducibility of dose responses on separate dates. In panels B-G, pairs of replicate cell viability measurements in the same cell line are plotted in scatterplots to demonstrate the high degree of similarity seen in measurements across all therapies tested **(B)**, pralatrexate **(C)**, bortezomib **(D)**, romidepsin **(E)**, gemcitabine **(F)**, and oxaliplatin **(G)**.

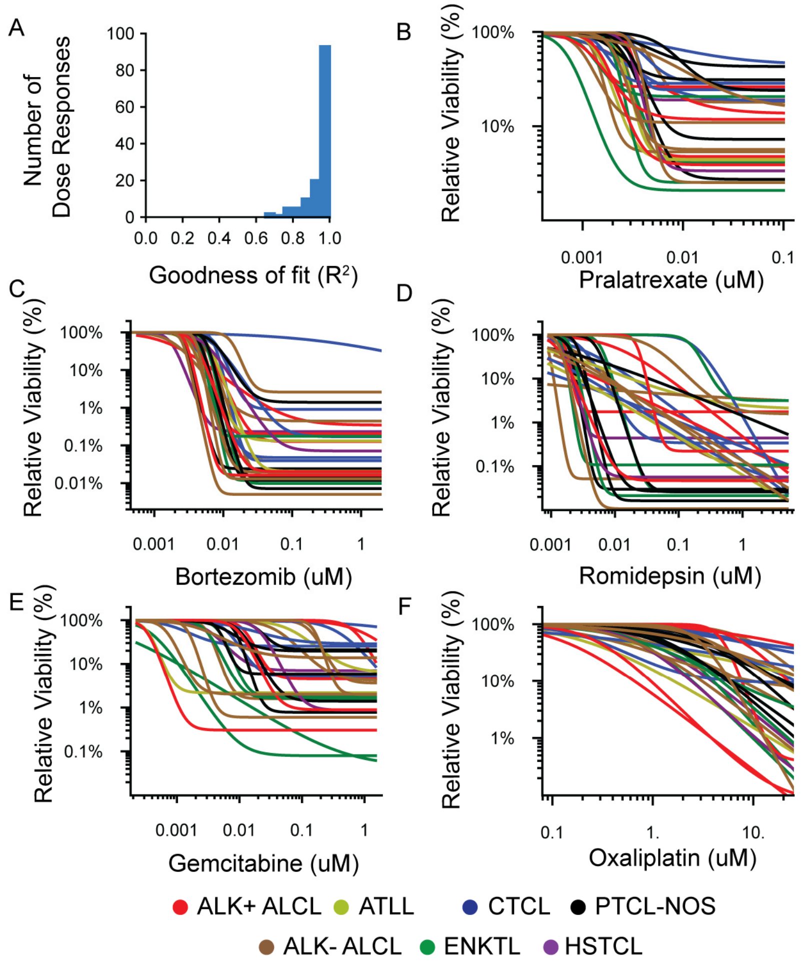


**Supplementary Figure 2. Immunophenotype-defined subgroups do not distinguish drug sensitivities for Mature T and NK cultures.** Dose response measurements were collected for 30 human T and NK lymphoma cultures with CellTiterGlo viability measurements relative to untreated controls after 72 hours of treatment. Decreasing Hill curves were fitted to a minimum of triplicate measurements. **A**. Goodness of Fit for 150 fitted curves shows greater than 75% (117/150) fits exceed an $R^2$ of 0.9. Fitted curves are presented with cultures categorized by immunophenotype for pralatrexate **(B)**, bortezomib **(C)**, romidepsin **(D)**, gemcitabine **(E)**, and oxaliplatin **(F)**. Generally cultures of the same subtype show a broad range of sensitivities to each therapy tested.

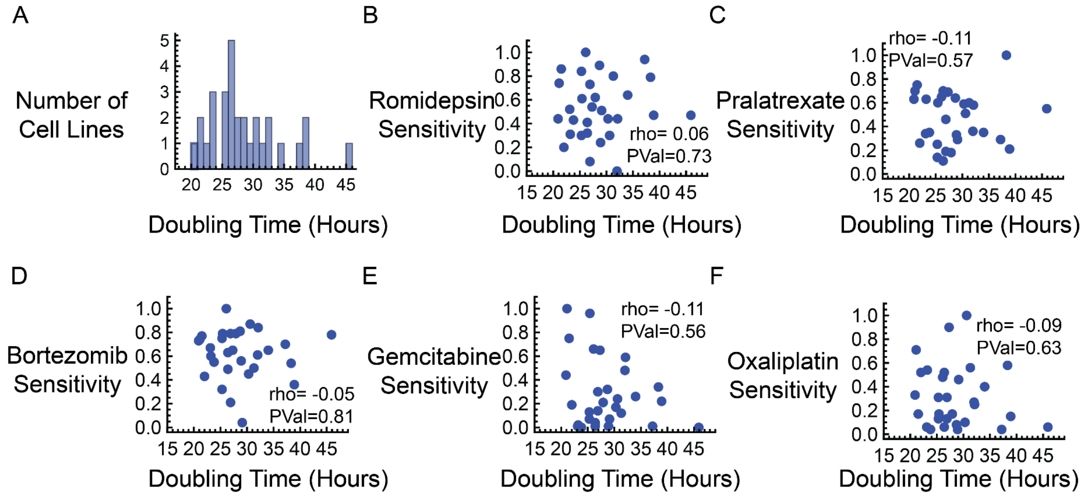


**Supplementary Figure 3. TCL Culture Growth Rate has no significant correlation with sensitivity to second line therapies.** Cell culture doubling times were measured by allowing each T or NK culture to grow in a minimum of 3 replicate wells of 12 well culture plates for a period of 72 hours in complete Human Plasma Like Medium. Live cell counts were measured in triplicate at the beginning and end of the growth period and used to calculate each culture's doubling time. **A.** Histogram of doubling times shows that the rate of growth for these T and NK cultures is largely consistent. The average time for a culture to double was 28.6 hours, with a standard deviation of 5.8 hours. Only two cultures exhibited doubling times greater than 1 standard deviation from the mean, the ALK- ALCL Culture SR-786 (45.9 hours), and the CTCL culture HuT-78 (38.9 hours). No significant correlation is calculated for doubling times and normalized sensitivity of cultures to (**B**) romidepsin, (**C**) pralatrexate, (**D**) bortezomib, (**E**) gemcitabine or (**F**) oxaliplatin. Spearman rank correlations and corresponding P values were calculated in Wolfram Mathematica.

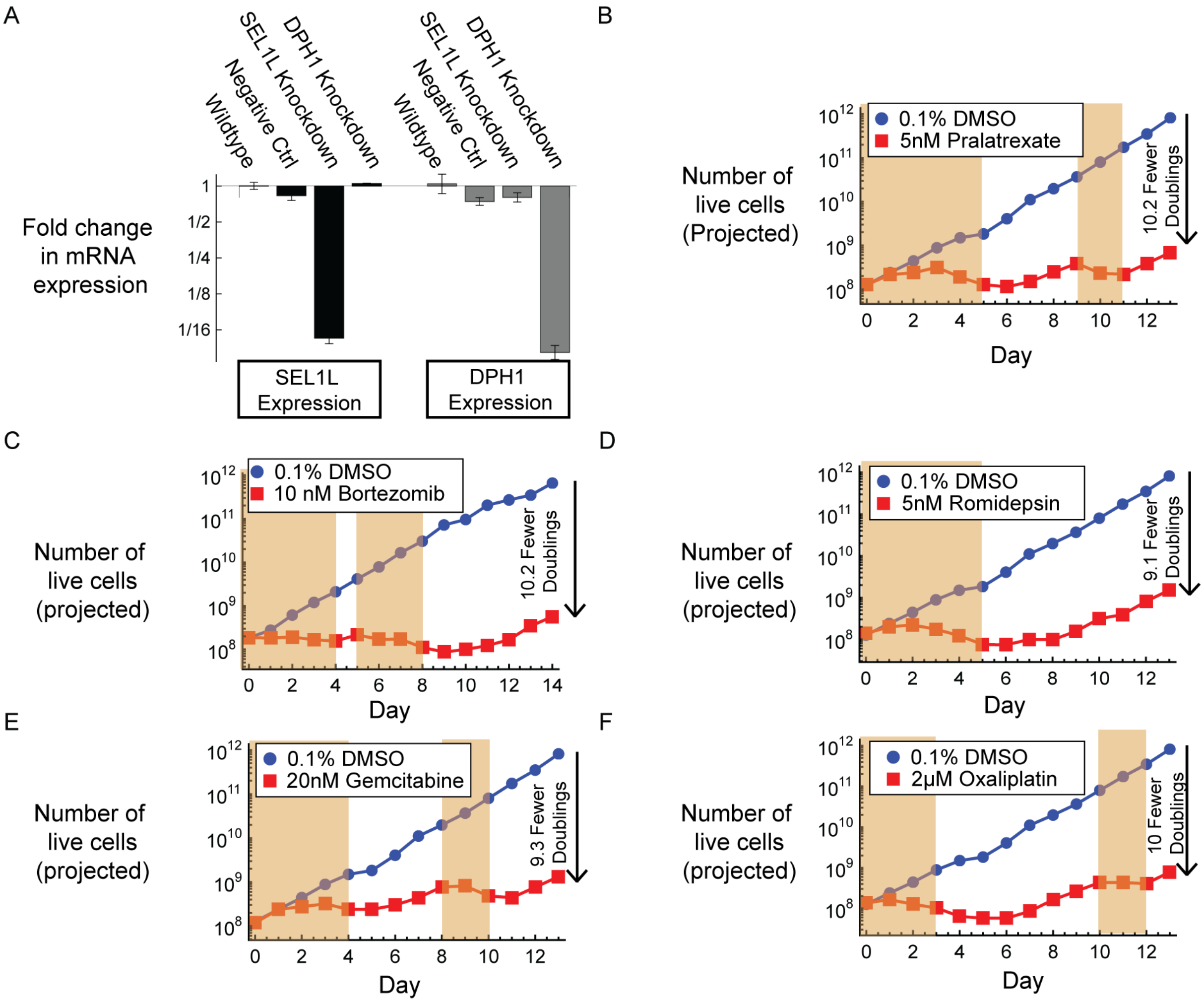


**Supplementary Figure 4. CRISPR Interference knockdown validation and growth projections for CRISPR Interference Screens with SU-DHL-1. A.** qPCR validation of CRISPR interference knockdown specificity in SU-DHL1 cells expression CRISPR-dCas9-KRAB construct. Knockdowns were validated with triplicate RNA extractions from single SU-DHL-1 cultures, SEL1L and DPH1 were randomly selected genes for knockdown validation with no specific relevance to the sensitivity of any second line therapy. **(B-F).** Fourteen day dosing schemes were followed with 5nM pralatrexate **(B)**, 0.01 nM Bortezomib **(C)**, 5nM romidepsin **(D)**, 20nM Gemcitabine **(E)**, and 1mM oxaliplatin **(F)**. Shaded regions on each plot indicate the days on which cultures were incubated with drugs at each concentration of interest. Dosing schemes were guided by daily measurements of growth and cell viability to enable 8-10 fold depletion of the drug treated culture as compared to a DMSO Control.

**Supplementary Table 2. Summary of Gene Knockdowns with significant phenotype scores in CRISPRi screens**

| Therapy | Number of genes above 10% FDR[a] | Maximum Mann-Whitney P Value of Phenotype Score | Common Pathway Associations Among CRISPR Hits[b] |
|---|---|---|---|
| Pralatrexate | 697 | 0.049 | Ribosome (MRPL27, RPL8, MRPL3, MRPS5, MRPL17, MRPL10, MRPL4, MRPL11)<br>Basal Transcription Factors (TAF4, CDK7, TAF5L, GTF2H4, MNAT1, GTF2H1, GTF2H2, GTF2E1)<br>Oxidative Phosphorylation (NDUFB4, NDUFB5, NDUFB10, COX17, ATP5F1B, ATP6V1A, ATP6V1B2, ATP5PO) |
| Bortezomib | 327 | 0.039 | Proteasome (PSMB3,4,5,6,7; PSMC1,2,3,4; PSMD1,2,3,4; PSME2)<br>Spinocerebellar Ataxia[c]<br>Huntington Disease[c] |
| Romidepsin | 142 | 0.029 | Basal Transcription Factors (MNAT1, GTF2E1, GTF2H1, GTF2H2, GTF2H2C, GTF2H3 GTF2H4)<br>Nucleotide Excision Repair (MNAT1, GTF2H1, GTF2H2, GTF2H2C, GTF2H3, GTF2H4)<br>Viral carcinogenesis (CCNA1, GTF2H4, CDK6, GTF2H1, GTF2H2, GTF2E1, BAK1, CDKN1A) |
| Gemcitabine | 503 | 0.048 | RNA transport (EIF3H, EIF2S2, NUP100, NUP188)<br>Cell Cycle (TP53, CDC23, E2F1, E2F3, CCNA2, CCNE1)<br>mRNA surveillance pathway (RNGTT, CCNT1, CCT2, KPNA3) |
| Oxaliplatin | 161 | 0.043 | Fanconi anemia DNA repair pathway (BRIP1, FANCM, FANCD2, FANCI, ATR)<br>p53 signaling pathway (TP53, BCL2L1, BAX)<br>Thyroid hormone signaling pathway (MED12, MED13L, KAT2A, HIF1A) |

[a] False Discovery Rate is determined based on the enrichment of pseudogenes, constructed by random draws from the distribution of phenotype scores measured for negative control guide RNAs

[b] Top Three KEGG pathways associated with CRISPR hits, identified using STRING Database 12.0 for all genes above False Discovery Rate of 10% for each therapy

[c] Many of the proteasome-related features (PSMB1, etc.) are associated with spinocerebellar ataxia and Huntington disease.

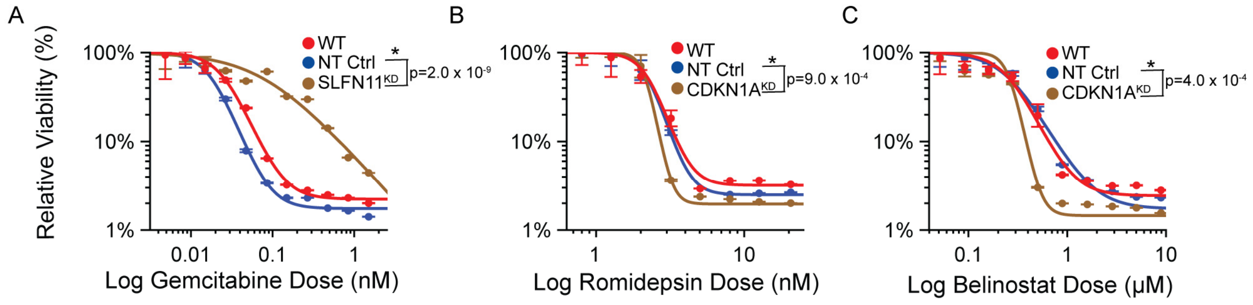


**Supplementary Figure 5. Validation of CRISPR interference knockdowns' impact on drug sensitivity of ALK+ ALCL culture.** Single gene knockdowns were performed with top performing guide RNAs from CRISPR interference screen. SU-DHL1 cultures were measured to be >90% expressing single guide RNAs against target genes of interest based on TagBFP expression measured with an Attune NxT Flow Cytometry System. **A**. Targeted knockdown of SLFN11 in SU-DHL-1 was observed to drive drug resistance over a range of doses greater than 0.01mM for Gemcitabine. **B and C**. CDKN1A knockdown increases sensitivity to Histone Deacetylase inhibitors romidepsin and belinostat. For romidepsin, the sensitization caused by CDKN1A knockdown was modest but significant (Student T-Test p-value $9.0 \times 10^{-4}$, comparing replicate IC90 values between SU-DHL1 cells expressing a non-targeting control guide and those expressing a guide to knockdown CDKN1A). Relatively greater impacts on belinostat sensitivity were measured (Student T-Test p-value $4.0 \times 10^{-4}$). Error bars reflect standard deviation of measurements from four replicate dose responses, with dot markers showing the average measured viability. P-values reflect t-tests comparing the dose required to kill 90% of cells from each replicate fit (n=4) between cultures with knockdown corresponding to target gene of interest versus cultures expressing a non-targeting control guide (NT-Ctrl).

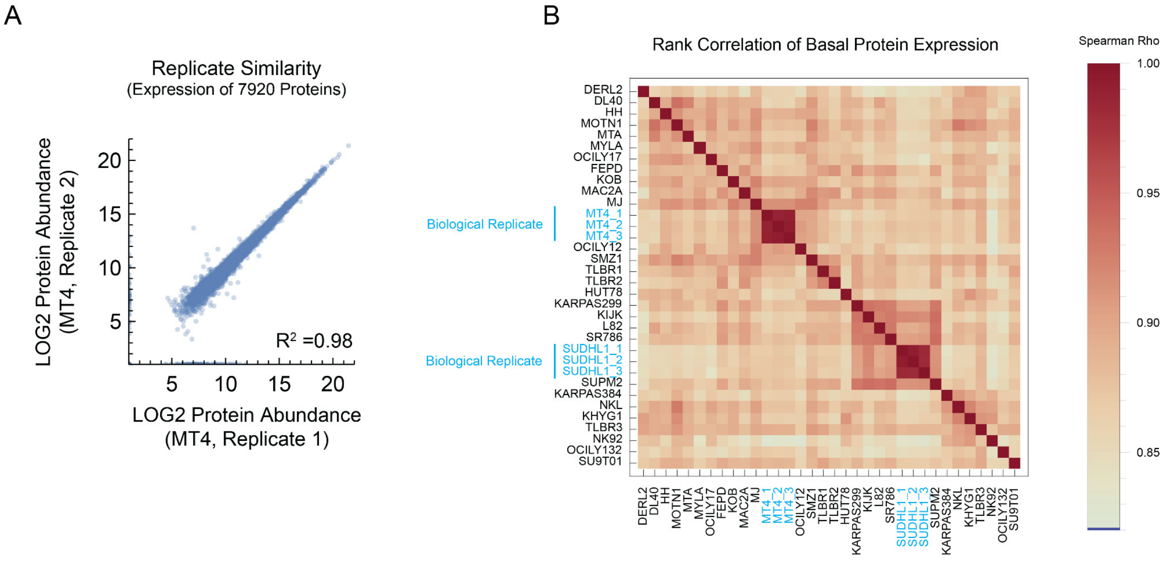


**Supplementary Figure 6. Proteomic profiles of 30 T and NK lymphoma cultures show a high degree of similarity.** We used LC/MS-MS with 20 micrograms of total protein from each of 30 T and NK lymphoma cell cultures to profile their basal protein expression. **A)** The correlation in basal protein expression measured between biologic replicates of the same parent culture is very high indicating reproducibility of LC/MS-MS proteomic profiling. **B)** Hierarchical clustering based on spearman rank correlation of protein expression across the whole panel of T and NK cultures reflects high overall similarity in proteomics for the whole panel of cultures with a minimum Spearman Rank Correlation of ~0.8. Triplicate biological replicates of the same cultures have the strongest degree of correlation.

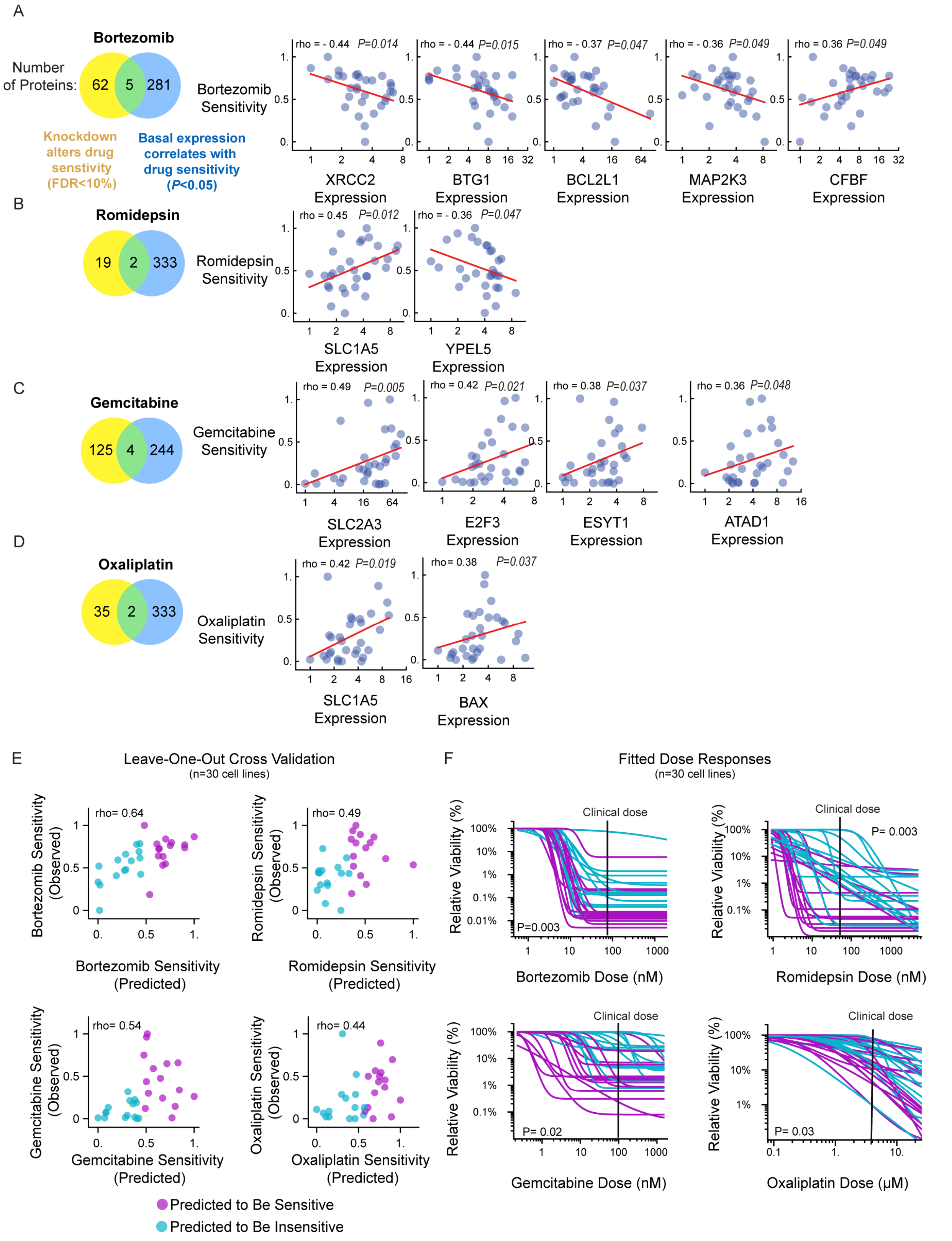


**Supplementary Figure 7. Identification of candidate biomarkers for additional second line therapies beyond pralatrexate.** (continued over page).

**Supplementary Figure 7. Identification of candidate biomarkers for additional second line therapies beyond pralatrexate. (A-D).** Venn diagrams demonstrating the identification of candidate biomarkers for bortezomib, romidepsin, gemcitabine, and oxaliplatin from hundreds of potential correlates and CRISPR interference screen hits. For each therapy of interest, the correlation between basal expression of candidate biomarkers and drug sensitivity is depicted with scatter plots and spearman rank correlation calculations. **E.** Leave-one-out cross validations were performed using linear regression between the sum of z scores of basal expression of candidate biomarkers and drug sensitivities of 30 T and NK Cell lines. For all therapies, the expression of these candidate biomarkers enables prediction of actual drug sensitivities with statistically significant correlation between the rank orders of predicted and actual area over the dose response curves. **F.** By stratifying cell lines into groups predicted to be sensitive (purple) or insensitive (bright blue), cell lines are distinguished into groups with significantly different observed sensitivities.

**Supplementary Table 3. Candidate biomarkers of drug sensitivity to second line therapies and literature support for their mechanistic relationships**

| Therapy | Gene | Effect of Knockdown | Literature Support | References |
|---|---|---|---|---|
| 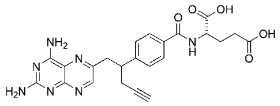 Pralatrexate | SLC19A1 | Resistance | SLC19A1 is the major folate transporter and it facilitates the transport of anti-folates including pralatrexate and methotrexate. Decreased SLC19A1 expression has been observed in pralatrexate-resistant cancer cells | (Hou and Matherly, 2014)[29] (Serova et al, 2011)[30] (Takimoto et al, 1996)[31] |
| | PEX5 | Resistance | Further investigation required | |
| | SLC1A5 | Resistance | Further investigation required | |
| 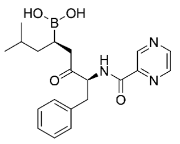 Bortezomib | BCL2L1 | Hypersensitivity | BCL2L1 is a key anti-apoptotic protein in the intrinsic apoptosis pathway, Mantle Cell Lymphoma cells that have evolved resistance to bortezomib exhibit overexpression of BCL2L1 | (Luanpitpong et al, 2022)[32] |
| | MAP2K3 | Hypersensitivity | MAP2K3 phosphorylates p38 for its activation, p38 inhibition has been shown in vitro to enhance bortezomib sensitivity in Multiple Myeloma cells | (Morales-Martinez and Vega, 2022)[33] (Hideshima et al, 2004)[34] |
| | BTG1 | Hypersensitivity | BTG1 is a tumor suppressor, low expression is generally associated with malignancy and chemoresistance. However, an independent genome wide siRNA screen identified BTG1 knockdown as synthetic lethal with bortezomib | (Kim et al, 2022)[35] (Chen et al, 2010)[36] |
| | CBFB | Resistance | Further investigation required | |
| | XRCC2 | Hypersensitivity | Further investigation required | |

(continued over page)

| | | | | |
|---|---|---|---|---|
| 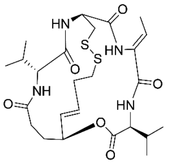<br>Romidepsin | SLC1A5 | Resistance | SLC1A5 is a transporter of glutamine, and glutamine metabolism inhibition has been shown to influence HDAC inhibitor sensitivity | (Okabe et al, 2024)[37] |
| | YPEL5 | Hypersensitivity | Further investigation required | |
| 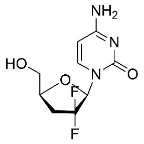<br>Gemcitabine | SLC2A3 | Resistance | Glucose Transporter 3 is known to transport capecitabine, and is thought to transport gemcitabine in breast and pancreatic cancer cells. Upregulation of SLC2A3 has been shown to promote gemcitabine sensitivity | (Chen et al, 2024)[38] |
| | E2F3 | Resistance | E2F3 is a transcription factor whose upregulation has been identified in the mediation of DNA-damage-induced apoptosis | (Martinez et al, 2009)[39] |
| | ESYT1 | Resistance | Further investigation required | |
| | ATAD1 | Resistance | Further investigation required | |
| 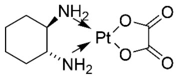<br>Oxaliplatin | BAX | Resistance | BAX is a proapoptotic factor in the intrinsic pathway of apoptosis, oxaliplatin sensitivity is influenced by BAX upregulation in cancer cell models | (Hayward et al, 2004)[40]<br>(Seo et al, 2022)[41] |
| | SLC1A5 | Resistance | SLC1A5 is a transporter of glutamine, and reducing glutamine metabolism has been shown to increase susceptibility to oxaliplatin in colorectal cancer models | (Lu et al, 2017)[42] |

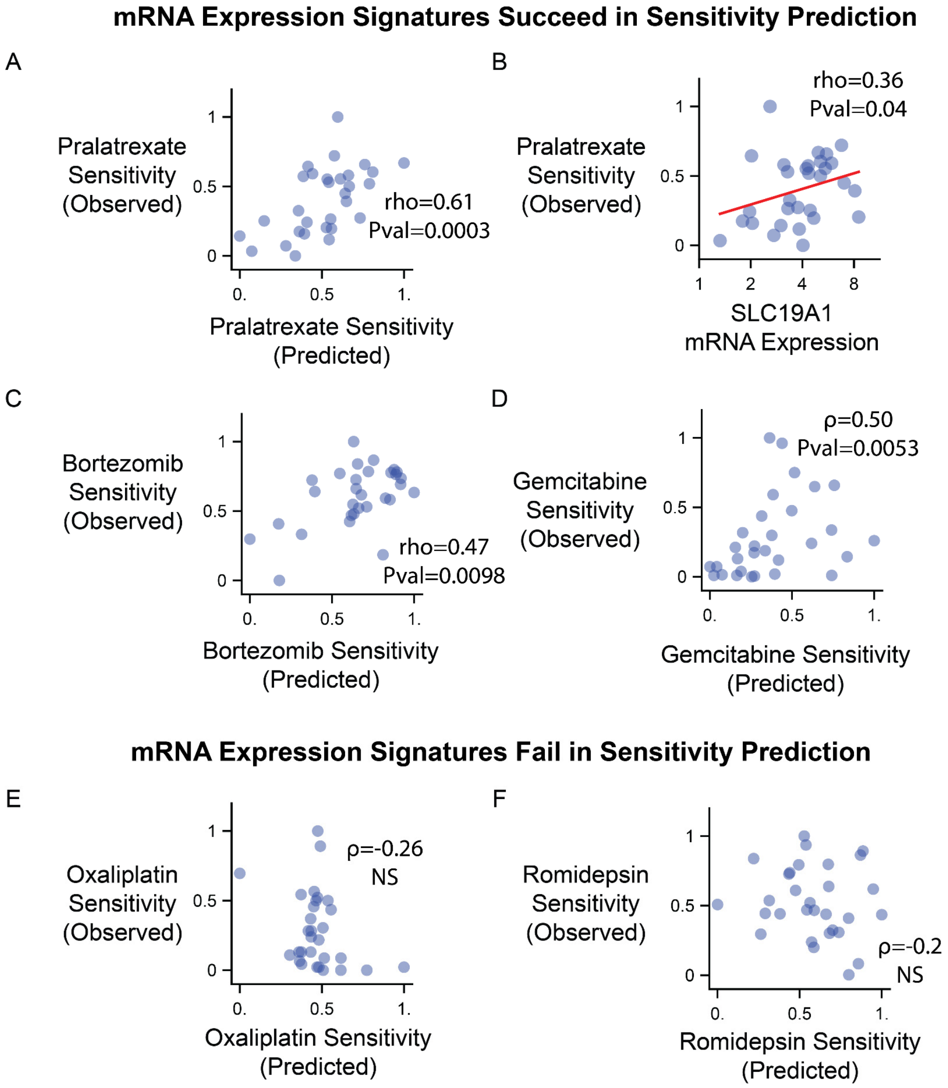


**Supplementary Figure 8. mRNA expression of candidate biomarkers is predictive of sensitivity for some but not all drugs.** In addition to performing proteomic profiling of T and NK cultures, we collected bulk RNA-seq measurements of basal gene expression for 30 T and NK lymphoma cultures. The mRNA transcript levels of the candidate biomarkers that we identified in this study can in some cases be used for the prediction of drug sensitivity as well as the protein expression differences. Generally sums of Z scores of mRNA transcript levels predict drug sensitivity with less accuracy by Leave-one-out cross validation than protein expression level, but predictions are still accurate for pralatrexate (A), bortezomib (C), gemcitabine (D). Panel B indicates the correlation between SLC19A1 mRNA expression and pralatrexate sensitivity. mRNA-based predictions of sensitivity were not accurate for oxaliplatin (E) or romidepsin (F).

**Supplementary Table 4. Parameter values for clinical trial simulations**

| Parameter | Symbol | Value | Units |
|---|---|---|---|
| Mean $Log_{10}$ (initial tumor population) | $\mu_p$ | 10.4 | $Log_{10}$ (cells) |
| Standard Deviation of $Log_{10}$ (initial tumor population) | $\sigma_p$ | 0.94 | $Log_{10}$ (cells) |
| Mean natural logarithm of tumor growth rate | $\mu_{growth}$ | –1.355 | Ln (week$^{-1}$) |
| Standard Deviation of natural logarithm of tumor growth rate | $\sigma_{growth}$ | 0.615 | Ln (week$^{-1}$) |
| Cell-to-cell heterogeneity in $Log_{10}$ (drug sensitivity) | $\sigma_{cell}$ | 0.20 | Dimensionless |
| Patient-to-patient heterogeneity in $Log_{10}$ (drug sensitivity) | $\sigma_{patient}$ | 1.1 | Dimensionless |
| Cell-to-cell correlation in $Log_{10}$ (drug sensitivity) | $\rho_{cell}$ | 0.21 | Dimensionless |
| Patient-to-patient correlation in $Log_{10}$ (drug sensitivity) | $\rho_{patient}$ | 0.44 | Dimensionless |
| Mean $Log_{10}$ (drug sensitivity) for drugs in CHOP | $\mu_{CHOP}$ | –0.60 | Dimensionless |
| Mean $Log_{10}$ (drug sensitivity) for Pralatrexate | $\mu_{pralatrexate}$ | –0.35 | Dimensionless |

The model structure is as described in Pomeroy & Palmer (2025) *Blood Cancer Discovery*. Each virtual patient's initial tumor population is sampled as $Log_{10}$(initial tumor population) ~ Normal($\mu_p$, $\sigma_p^2$). Each virtual patient's tumor growth rate is sampled as Ln(growth rate, in weeks$^{-1}$) ~ Normal($\mu_{growth}$, $\sigma_{growth}^2$). To illustrate, the mean Ln(growth rate) of –1.355 corresponds to growth rate constant = exp(–1.355) = 0.26 weeks$^{-1}$, or equivalently, a doubling time of Ln(2) / 0.26 = 2.7 weeks. Each virtual patient's $Log_{10}$(drug sensitivities), as a 5-element vector $\boldsymbol{x}$ for drugs [C, H, O, P, pralatrexate], is sampled as $\boldsymbol{x}$ ~ Multinormal([$\mu_{CHOP}$, $\mu_{CHOP}$, $\mu_{CHOP}$, $\mu_{CHOP}$, $\mu_{pralatrexate}$] , $\boldsymbol{\Sigma}_{patients}$) where $\boldsymbol{\Sigma}_{patients}$ is a covariance matrix of patient-to-patient variation with diagonals elements $\sigma_{patient}^2$ and off-diagonal elements $\sigma_{patient}^2\ \rho_{patient}$. Each virtual patient's tumor population is then modelled as a population of cells with a distribution of drug sensitivities modelled as Multinormal($\boldsymbol{x}$, $\boldsymbol{\Sigma}_{cells}$) where $\boldsymbol{\Sigma}_{cells}$ is a covariance matrix of cell-to-cell variation with diagonals elements $\sigma_{cell}^2$ and off-diagonal elements $\sigma_{cell}^2\ \rho_{cell}$. At each coordinate of the distribution the fraction of tumor cells surviving a cumulative dose **D** of chemotherapy (expressed as a vector of doses of C, H, O, P, pralatrexate) is exp(–$10^{\boldsymbol{x}}$ **D**). Doses per cycle are 1 by definition; differences in drug effectiveness are accounted by the mean drug sensitivity values. CHOP consists of 6 cycles, each of 2 weeks duration, composed of dose [1, 1, 1, 1, 0] at the start of the first week. Pralatrexate monotherapy for R/R PTCL consists of 7-week cycles composed of 6 weekly doses of [0, 0, 0, 0, 1] and a seventh week off-therapy. COP-Pralatrexate consists of 6 cycles, each of 2 weeks duration, with dose [1, 0, 1, 1, 1] (i.e. doxorubicin is replaced by pralatrexate) at the start of the first week and dose [0, 0, 0, 0, 1] (i.e. pralatrexate only) at the start of the second week.
